# The unique microbial diversity, community composition and networks of Pacific Islander endocervical and vaginal microbiomes in the presence or absence of *Chlamydia trachomatis* infection using metagenomic shotgun sequencing

**DOI:** 10.1101/2023.06.23.546186

**Authors:** Sankhya Bommana, Yi-Juan Hu, Mike Kama, Ruohong Wang, Reshma Kodimerla, Kenan Jijakli, Timothy D. Read, Deborah Dean

## Abstract

**Background:** Pacific Islanders are a vulnerable population with a high prevalence of *Chlamydia trachomatis* (*Ct*) sexually transmitted infections (STIs) and remain underrepresented in research. Here, 258 vaginal and 92 paired endocervical samples from women of diverse ethnicities in Fiji were evaluated using metagenomics to characterize microbial relative abundance, composition and networks including associations with *Neisseria gonorrhoeae*, human papilloma virus (HPV), *Mycoplasma genitalium*, *Candida* and bacterial vaginosis (BV).

**Results:** Pacific Islander ethnicities and age <25 years were significantly associated with *Ct* infection. Using VALENCIA, a sub-community state type (subCST) classifier, 93 (36.04%) vaginal and 38 (41.30%) endocervical microbiomes did not match reference subCSTs. Four unique subCSTs were developed to better classify Pacific Islander microbiomes: IV-D0, dominated by *Gardnerella vaginalis*; IV-D1, high/moderate *G. vaginalis* relative abundance with *Prevotella* spp.; IV-D2, high/moderate *G. vaginalis* with *Lactobacillus iners*; and IV-E, moderate *Prevotella* spp*. Ct* positive endocervical and paired vaginal microbiomes were significantly more likely to have differential species relative abundance (81.58%) than *Ct* negative pairs (35.84%; AOR: 7.93; 95% CI: 2.93-21.93; *P*<0.0001). A significantly higher alpha diversity was found for iTaukei ethnicity, all subCST-IV types, BV, and *Ct* for vaginal microbiomes. For endocervical microbiomes, higher diversity was significant for subCST-IV-A, subCST-IV-D1, and subCST-IV-E, and high-risk HPV types. Overall, there was a significantly higher diversity for the endocervix in paired microbiomes. Vaginal microbiomes showed significant divergence in community composition as above and for *Candida.* Endocervical composition varied significantly by subCST type and *Ct* status. Gut and BV-associated bacterial clusters were present in *Ct* positive and negative paired endocervical and vaginal microbiome networks but were smaller with fewer bacterial and no *Lactobacillus* spp. interactions in *Ct*-infected endocervical networks where *G. vaginalis* generated polymicrobial biofilms along with *Ct* likely influence pathogenicity.

**Conclusions:** Fijian endocervical and vaginal microbiomes represent divergent microbial abundance and compositions, especially for Pacific Islander ethnicities, with distinct subCSTs compared to other global populations. The higher microbial diversity of the endocervix with prevalent *G. vaginalis*, *L. iners* and *Prevotella* spp. suggest that these microbiomes/networks may predispose to and/or promote chlamydial and HPV pathogenesis. Prospective studies are needed to further define causal associations to develop successful interventions.

## Background

*Chlamydia trachomatis* (*Ct*), a bacterial pathogen restricted to humans, is the most common cause of sexually transmitted infections (STIs) worldwide. The World Health Organization (WHO) estimates that over 130 million global cases occur each year with a steady increase since the 1990s [1,2]. The distribution of these infections is greatest in Africa and the Western Pacific Region (WPR) where the WPR has the highest incidence at 61 million cases [1]. The prevalence of *Ct* STIs ranges from about 30% to 44% among teens and young adults in the Pacific Island Countries and Territories (PICT) of the WPR [3–6]. Here, the combination of asymptomatic, untreated infection and syndromic management, which is based on signs and symptoms as a proxy for STI diagnostics, can lead to life-threatening ectopic pregnancy, infertility, and an increased risk of cervical cancer and HIV [7–10]. Indeed, PICTs have the third highest prevalence of infertility in the world today [11].

The composition of the vaginal microbiome is influenced by resident microbes but also bacterial vaginosis (BV) and STIs including *Ct, Neisseria gonorrhoeae* (*Ng*), *Trichomonas vaginalis* (*Tv*), *Mycoplasma genitalium* (*Mg*), HIV and Human Papilloma Virus (HPV) [12–14]. Community state types (CSTs) were developed to classify vaginal microbiomes that include CST-I, CST-II, CST-III, CST IV and CST-V [15]. CSTs I-III and V are dominated by *Lactobacillus crispatus*, *Lactobacillus gasseri*, *Lactobacillus iners* or *Lactobacillus jensenni*, respectively, that maintain vaginal homeostasis by secreting lactic acid, bacteriocins, hydrogen peroxide, and antimicrobial compounds to inhibit the growth of anaerobic bacteria and protect against BV and STIs. CST-IV, however, is deficient in *Lactobacillus* spp. and comprised of anaerobic bacteria largely from the *Prevotella*, *Dialister*, *Atopobium*, *Gardnerella*, *Megasphaera*, *Peptoniphilus*, *Sneathia*, *Eggerthella*, *Aerococcus*, *Finegoldia* and *Mobiluncus* genera. Change from a *Lactobacillus* dominant to a diverse polymicrobial microbiome can increase the risk of acquiring *Ct* and other STIs [16–18]. Vaginal microbiome data from European, Asian, African and Hispanic populations have shown that distinct CSTs are found among these racial, ethnic and geographic groups regardless of STI status [19–23]. However, data from other ethnic and racial populations such as indigenous Australian, Native American and Pacific Islander populations are lacking.

In an effort to provide insights into the epidemiology, structure, and function of the human vaginal microbiome, the vaginal non-redundant gene catalog (VIRGO) [24] and subCST classifier VALENCIA [25] were developed in 2020. The former was constructed from 264 vaginal metagenome shotgun sequences (MSS) and 308 draft genomes of urogenital bacterial isolates primarily obtained from North American women and tested using 91 vaginal metagenomes from North American, African and Chinese women. These data provided over 95% vaginal microbiome coverage applicable to these populations. VALENCIA classifies microbiomes into 13 subCSTs based on their similarity to a set of reference centroids built using a comprehensive set of 13,160 16S rRNA sequencing-based taxonomic profiles from 1975 North American Black, White, Hispanic and Asian women. It was validated using datasets of reproductive age eastern and southern African women and adolescent girls, and ethnically and geographically diverse samples of postmenopausal women, although the number of women in each dataset was not provided.

We recently implemented VIRGO v1.0 and VALENCIA v1.0 on paired endocervical, vaginal and rectal MSS of Pacific Islanders in Fiji as a pilot study, age-matching five women with *Ct* to those without *Ct* STIs [26]. VALENCIA v1.0 could not accurately classify 4 of 10 vaginal and 4 of 10 endocervical microbiomes. Furthermore, because VIRGO v1.0 does not contain viral, fungal or protozoan pathogen sequences, such as HPV, *Candida* and *T. vaginalis,* respectively, our analyses for these pathogens was limited. We found that the microbiomes were largely dominated by species other than *Lactobacillus* regardless of STI status and, while the endocervical and vaginal microbiomes were closely related, they were distinct in taxonomic composition, subCSTs and metabolic function. However, our pilot study was limited in sample size, and only ethnic iTaukei Pacific Islanders were studied.

There is an obvious need to expand MSS studies of endocervical and vaginal microbiomes from diverse ethnic and global populations to better understand the microbial diversity, compositions and networks that drive pathogenic microbiomes. Building on these existing knowledge gaps, we expanded our dataset to 258 vaginal and 92 paired endocervical samples using MSS to characterize vaginal and endocervical microbial relative abundance, composition and network interactions for the diversity of ethnicities among women residing in Fiji. We also explored differences between paired endocervical and vaginal microbiomes with and without the presence of *Ct*, other STIs, *Candida* and BV.

## Results

### Characteristics of study subjects, infection prevalence and quality statistics of endocervical and vaginal metagenomic shotgun sequences (MSS)

Women 18 to 40 years of age attending the Fiji Ministry of Health and Medical Services (MoHMS) health centers and outreach clinics were enrolled as part of a parent study as previously described [3]. The ethnic distribution in Fiji is approximately 57% iTaukei or indigenous Fijians followed by 37% Indo-Fijians [27,28]. Other ethnic groups include Chinese, European, mixed race, other Pacific Islanders as well as expatriates of various nationalities. Other Pacific Islanders comprise mostly IKiribati, Banabans, Tuvaluans, Tongans, Samoans and Wallisians [28].

From the parent study, 258 vaginal samples, of which 92 had paired endocervical samples, were available to the current study. Data on participants’ age, ethnicity, and STI status, including *Ct*, *Ng, Tv* and HPV, and Candida and BV were provided (**Supplementary Table 1)**. Age under 25 years and Pacific Islander ethnicities were significantly associated with *Ct* infection (**Table 1**).

**Table 1.**
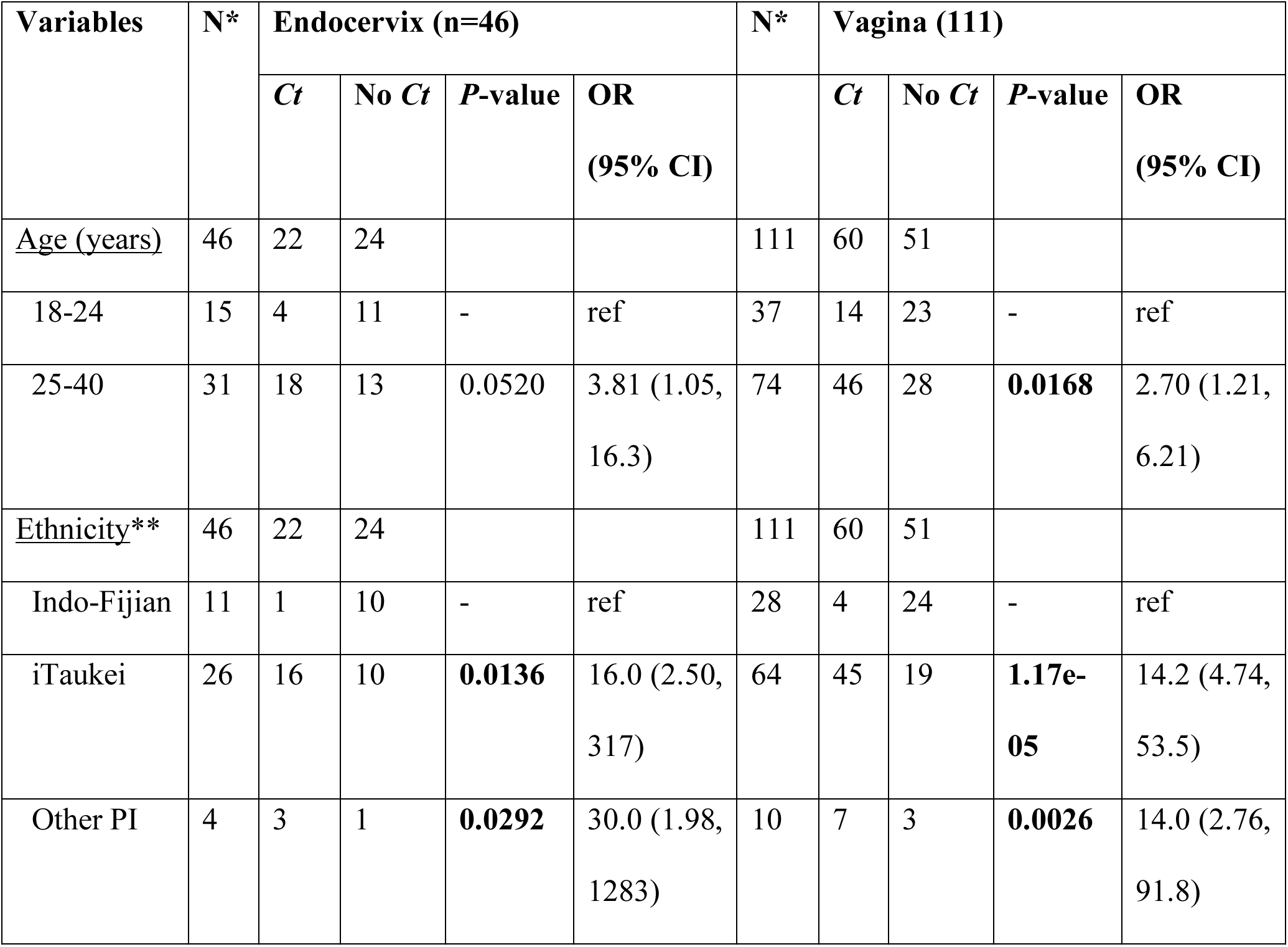
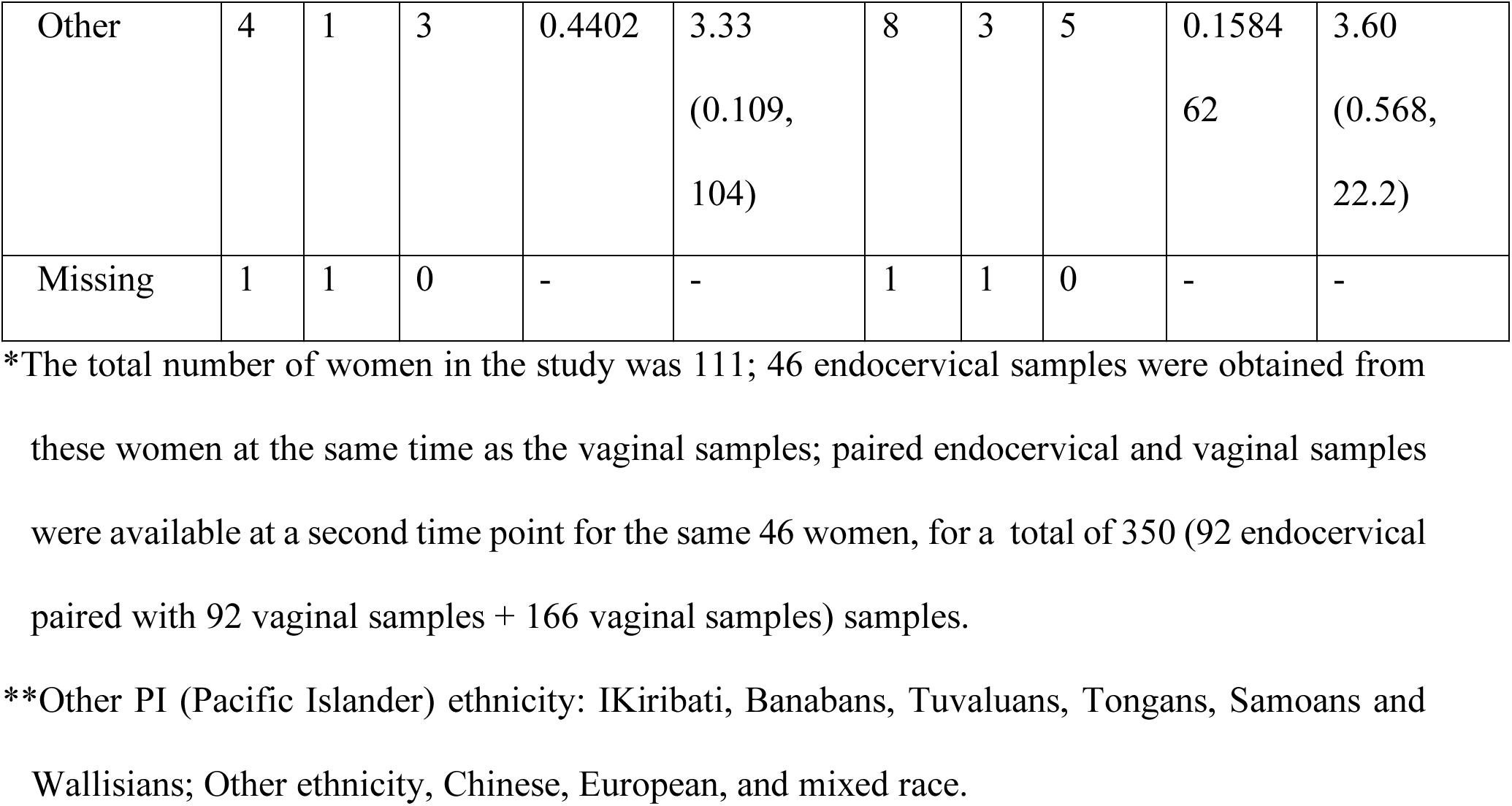
Bivariate analysis for age group and ethnicity associated with *C. trachomatis* infection for study participants.

Results for BV using Amsel criteria [29] and for *Tv* and *Candida*—detected by wet prep on vaginal samples (provided by the parent study) and MetaPhlAn v3.0 (this study) for both the vagina and endocervix—are also shown in **Supplementary Table 1**. In the multivariate logistic regression model, there was a significant association with *Ct* infection for *Ng* in both the endocervix and vagina, and with high-risk (hr)HPV, *Ng* and BV only in the vagina (**Table 2**).

**Table 2.** Bivariate analysis and multivariate model for sexually transmitted infections, including human papilloma virus (HPV), *M. genitalium*, *N. gonorrhoeae* and *T. vaginalis,* in addition to bacterial vaginosis (BV) and their association with *C. trachomatis* infection for endocervical and vaginal samples.

| Variables | N* | Endocervix (n=92) |  |  |  |  |  | N* | Vagina (n=258) |  |  |  |  |  |
| --- | --- | --- | --- | --- | --- | --- | --- | --- | --- | --- | --- | --- | --- | --- |
|  |  | Ct | No<br>Ct | P-value | OR<br>(95% CI) | P-value | AOR<br>(95% CI) |  | Ct | No<br>Ct | P-value | OR<br>(95% CI) | P-value | AOR<br>(95% CI) |
| <u>HPV**</u> | 92 | 40 | 52 |  |  |  |  | 258 | 99 | 159 |  |  |  |  |
| No infection | 38 | 13 | 25 | - | ref | - | - | 101 | 33 | 68 | - | ref | - | ref |
| lrHPV | 29 | 15 | 14 | 0.152 | 2.06<br>(0.771,<br>5.64) | - | - | 116 | 45 | 71 | 0.349 | 1.31<br>(0.748,<br>2.29) | - | - |
| hrHPV | 25 | 12 | 13 | 0.276 | 1.78<br>(0.633,<br>5.05) | - | - | 41 | 21 | 20 | **0.0410** | 2.16<br>(1.03,4.5<br>7) | **0.0329** | 2.45 (1.08,<br>5.66) |
| Missing | 0 | 0 | 0 | - | - | - | - | 0 | 0 | 0 | - | - | - | - |
| <u>M. genitalium</u> | 92 | 40 | 52 |  |  |  |  | 258 | 99 | 159 |  |  |  |  |
| Absent | 85 | 34 | 51 | - | ref | - | ref | 249 | 95 | 154 | - | ref | - | - |
| Present | 7 | 6 | 1 | **0.0463** | 9.00<br>(1.45,<br>174) | 0.0572 | 10.7 (1.35,<br>255) | 9 | 4 | 5 | 0.7040 | 1.30<br>(0.314,<br>5.02) | - | - |
| Missing | 0 | 0 | 0 | - | - | - | - | 0 | 0 | 0 | - | - | - | - |
| *N. gonorrhoeae* | 92 | 40 | 52 |  |  |  |  | 258 | 99 | 159 |  |  |  |  |
| Absent | 53 | 15 | 38 | - | ref | - | ref | 197 | 67 | 130 | - | ref | - | ref |
| Present | 39 | 25 | 14 | **0.0008** | 4.52<br>(1.90,<br>11.3) | **0.0132** | 3.49 (1.32,<br>9.70) | 61 | 32 | 29 | **0.0104** | 2.14<br>(1.20,<br>3.85) | **0.0410** | 1.94 (1.03,<br>3.68) |
| Missing | 0 | 0 | 0 | - | - | - | - | 0 | 0 | 0 | - | - | - | - |
| *T. vaginalis* | 92 | 40 | 52 |  |  |  |  | 258 | 99 | 159 |  |  |  |  |
| Absent | 86 | 37 | 49 | - | ref | - | - | 232 | 84 | 148 | - | ref | - | - |
| Present | 6 | 3 | 3 | 0.740 | 1.32<br>(0.234,<br>7.51) | - | - | 26 | 15 | 11 | **0.0368** | 2.40<br>(1.06,<br>5.60) | 0.282 | 1.60<br>(0.685,<br>3.85) |
| Missing | 0 | 0 | 0 | - | - | - | - | 0 | 0 | 0 | - | - |  |  |
| <u>Bacterial</u><br><u>Vaginosis</u> |  |  |  |  |  |  |  | 258 | 99 | 159 |  |  |  |  |
| Absent | - | - | - | - | - | - | - | 174 | 58 | 116 | - | ref | - | ref |
| Present | - | - | - | - | - | - | - | 39 | 24 | 15 | **0.00150** | 3.20<br>(1.58,<br>6.68) | **0.0153** | 2.64 (1.22,<br>5.93) |
| Missing | - | - | - | - | - | - | - | 45 | 17 | 28 | - | - | - | - |

NOTE: OR, Odds Ratio; AOR, Adjusted OR,
\*The total number of women in the study was 111; 46 endocervical samples were obtained from these women at the same time as the vaginal samples; paired endocervical and vaginal samples were available at a second time point for the same 46 women, for a total of 350 (258 + 92) samples for HPV, *M. genitalium*, *T. vaginalis*, and *N. gonorrhoeae* associations; these second time point vaginal samples were also used in the BV analyses (212 + 46). Note that BV cannot be diagnosed in the endocervix.
\*\*hrHPV, high-risk HPV; lrHPV, low-risk HPV

The 258 vaginal and 92 endocervical metagenomes yielded a total of ∼26 billion and 9.5 billion raw reads of which 22.8 billion (87.36%) and 8.76 billion (91.4%) were identified as human contamination, respectively (**Supplementary Table 2)**.

Based on VIRGO, *Mg* reads were detected in nine (3.5%) of 258 vaginal and seven (7.6%) of 92 endocervical metagenomes with a cut off of > 1 RPM (see Methods; **Supplementary Table 3**). HPV was identified by HPViewer [30] (**Supplementary Table 3**) in 157 (60.9%) of 258 vaginal and 54 (58.7%) of 92 endocervical samples; high-risk (hr)HPV types were found in 92 (35.66%) and 38 (41.30%) samples, respectively. However, not all endocervical and vaginal paired samples had the same HPV types. The most common low-risk (lr)HPV types in all samples were HPV 57 (n = 45; 25 vaginal and 20 endocervical) followed by HPV 62 (n = 42; 32 vaginal and 10 endocervical) and HPV 90 (n = 40; 26 vaginal and 14 endocervical). The most common hrHPV types were HPV 52 (n = 35; 24 vaginal and 11 endocervical) and 39 (n = 30; 17 vaginal and 13 endocervical). hrHPV types 16 and 18—responsible for most HPV-related cancers—were found in 24 (9.30%) of 258 vaginal and 11 (11.96%) of 92 endocervical samples and 18 (6.98%) vaginal and 21 (22.83%) endocervical samples, respectively. For *Tv*, six samples were positive by MSS alone in the endocervix and 22 in the vagina—10 by wet prep alone, 4 by both wet prep and MSS, and 8 by MSS alone (**Supplementary Table 1**). Few samples were positive for *Candida* in the vagina and none in the endocervix (**Supplementary Table 3**).

### Expansion of existing CST classification system to include new subCSTs unique to endocervical and vaginal microbiomes of Pacific Islanders

The vaginal and endocervical microbiomes resolved into the five CSTs described by Ravel *et al*. [15] and 13 subCSTs based on a nearest-centroid algorithm as defined by VALENCIA [25]. The algorithm assigns a similarity score of 0-1 between the microbial profile of the sample to one of the 13 reference centroids. In our previous [26] and current study, several samples had low similarity scores (<0.1-0.25), but also scores above 0.4 with a range of 0.019-0.808 where there was a discrepancy between the species abundance of the microbiomes and the archetypal profile of the closest reference centroid. We, therefore, manually compared the microbial relative abundance profiles of each vaginal and endocervical microbiome with that of the assigned subCST, re-classifying samples regardless of their similarity score.

The relative abundance profiles for 93 (36.04%) of the 258 vaginal and 38 (41.3%) of the 92 endocervical microbiomes did not match their assigned subCST (**Supplementary Table 4)**. These profiles indicated that they could be grouped into four new subCST categories within CST IV: IV-D0, dominated by *G. vaginalis* (n=51); IV-D1, a high to moderate relative abundance of *G. vaginalis* with *Prevotella* spp. (n=18); IV-D2, a high to moderate relative abundance of *G. vaginalis* with *L. iners* (n=18); and IV-E, moderate *Prevotella* spp. with other species (n=25) (**Figure 1**, orange highlights). These new reference subCSTs were constructed by averaging the relative abundance of each microbiota for all microbiomes (n=131) fitting the new subCST designation (**Supplementary Table 5**).

**Figure 1.**
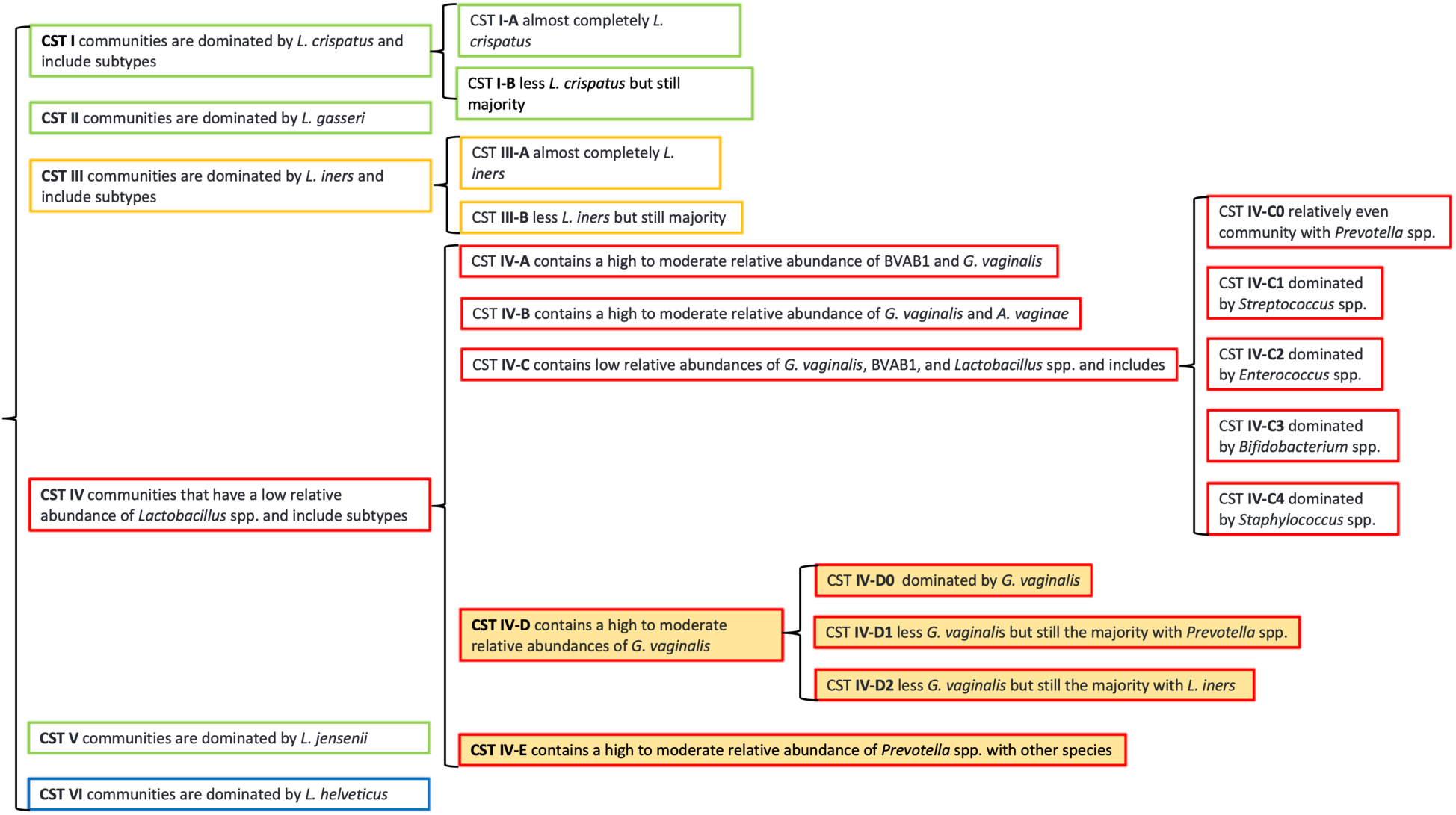
Classification of the endocervical and vaginal community state types (CSTs) by VALENCIA with inclusion of four new subCSTs (IV-D0, IV-D1, IV-D2 and IV-E) identified among Pacific Islanders residing in Fiji. The new subCSTs (denoted in orange background) were constructed by averaging the relative abundance of all microbiota for the samples that fit the new subCST designation (n=112; see Methods). A new CST VI suggested by Li *et al*. [31] is also included in the Figure (blue outlined box).

Microbiomes that we classified as subCST IV-D0, IV-D1 and IV-D2 were originally assigned CST IV-B by VALENCIA (**Figure 2**). While CST IV-B was defined as a high to moderate relative abundance of *G. vaginalis* (33.75%) and *A. vaginae* (12.16%), the new subgroups were different in composition. SubCST IV-D0 had a much higher relative abundance of *G. vaginalis* at 85.15% and much lower level of *A. vaginae* at 2.46%. SubCST IV-D1 had a higher relative abundance of *G. vaginalis* (52.72%) with a higher abundance of *Prevotella* spp. (20%), and subIV-D2 had a higher relative abundance of *G. vaginalis* (61.84%) with a higher abundance of *L. iners* (23.2%) compared to CST IV-B (**Figure 2**).

**Figure 2.**
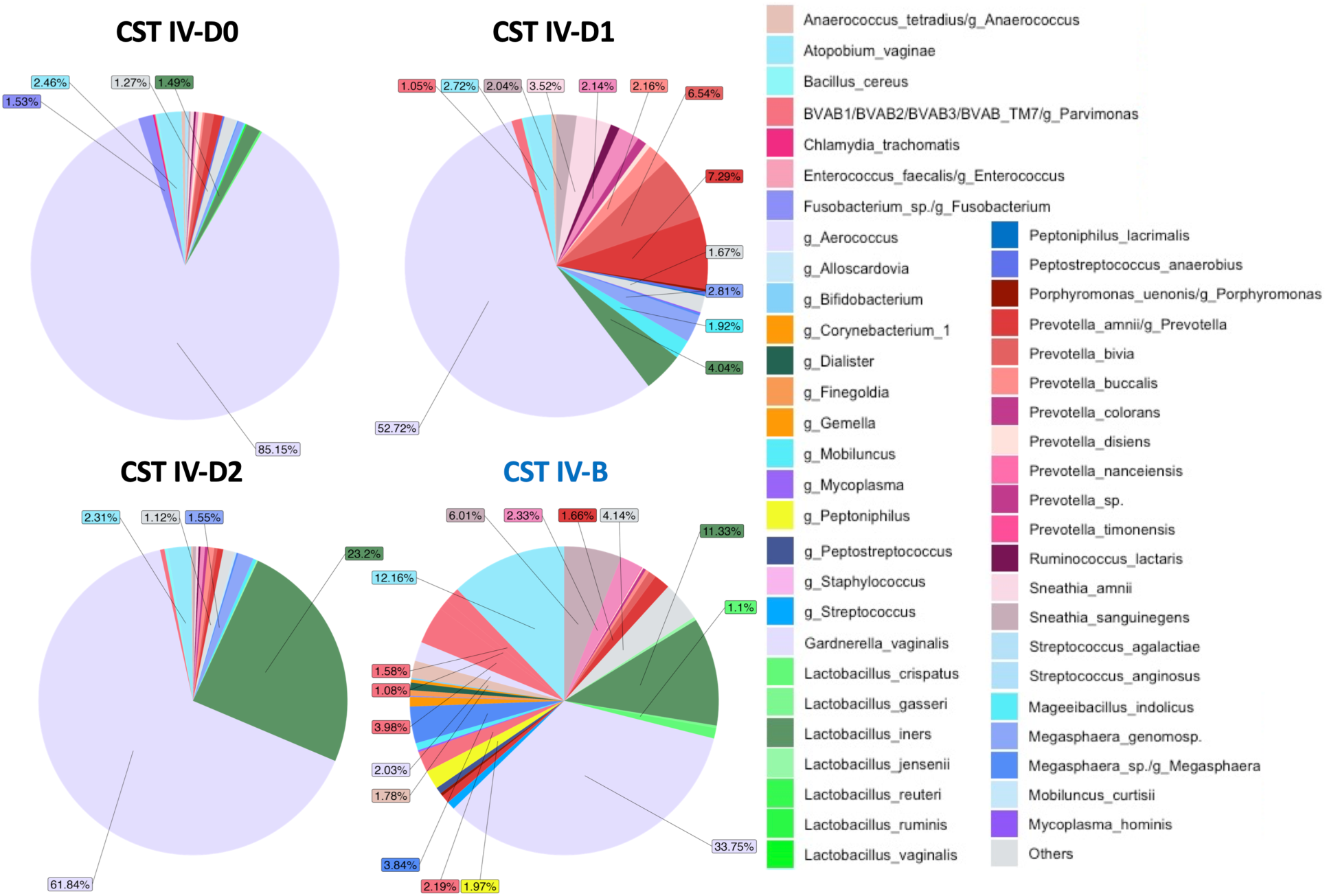
Pie chart representation of unique sub-community state types (subCSTs) IV-D0, IV-D1, and IV-D2 developed for endocervical and vaginal microbiomes from the Fiji metagenome database output from VIRGO. The microbiomes were originally assigned to subCST IV-B (right lower pie graph) using the VALENCIA classifier but were manually re-classified to the new subCSTs based on their relative abundance profiles that were not a good match with IV-B (see Methods). The new subCSTs were defined as IV-D0, dominated by *G. vaginalis;* IV-D1, less *G. vaginalis* but still the majority with *Prevotella* spp.; and IV-D2, less *G. vaginalis* but still the majority with *L. iners*. Relative abundance data were used to plot the pie charts of the new CSTs (**Supplementary Table 5**).

Samples that were originally assigned as CST III-B, IV-A, IV-B, IV-C0, IV-C1 and IV-C4 by VALENCIA were re-classified as subCST IV-E (**Figure 3**). CST IV-E, which was distinct from other CSTs, consisted of an approximate 50% relative abundance of *Prevotella* spp. and 12.29% for *G. vaginalis*.

**Figure 3.**
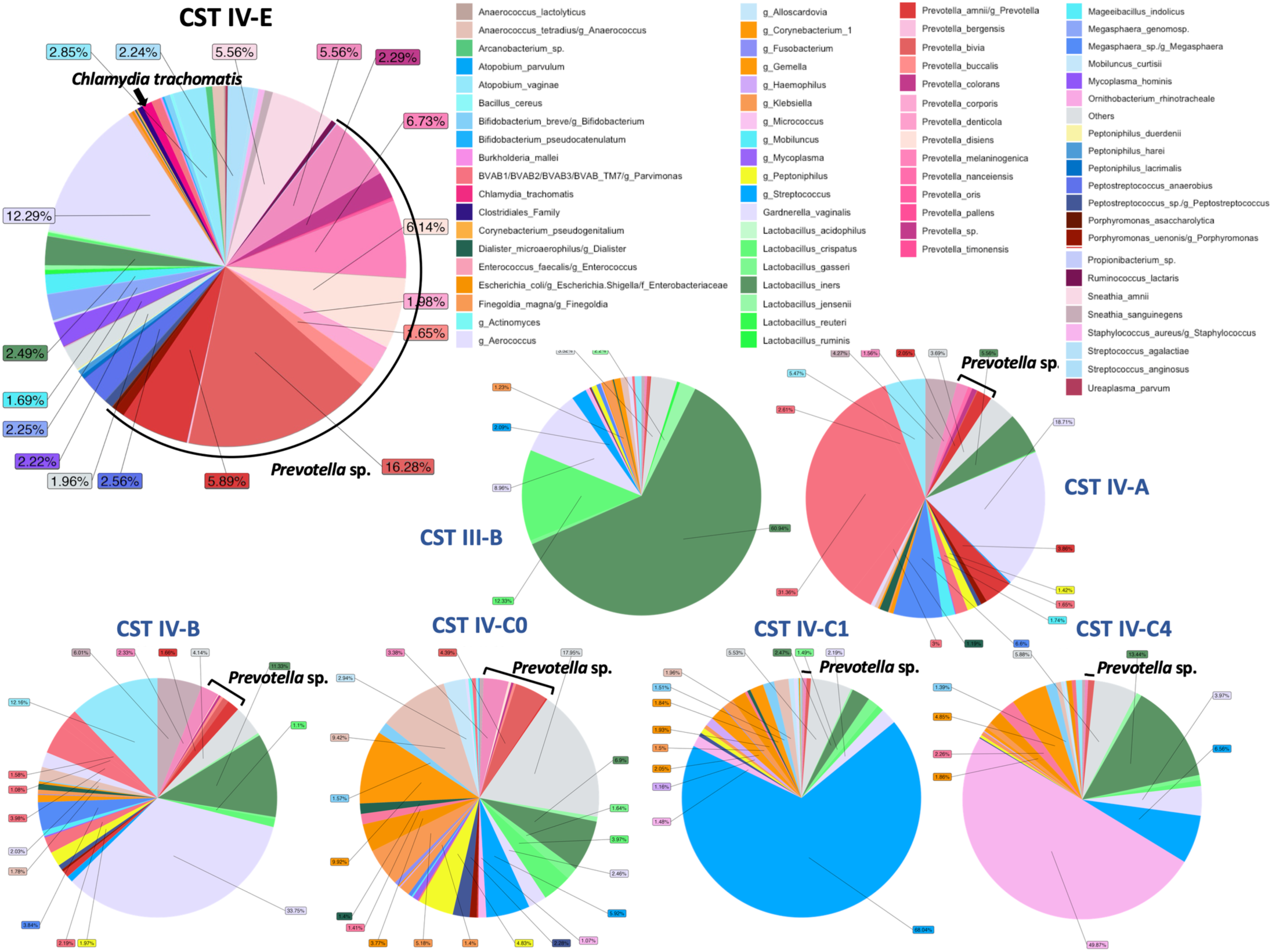
Pie chart representation of the unique sub-community state type (subCST) IV-E (upper left) developed for endocervical and vaginal microbiomes from the Fiji metagenome database output from VIRGO. The microbiomes were originally assigned to subCSTs III-B, IV-A, IV-B, IV-C0, IV-C1 and IV-C4 using the VALENCIA classifier (denoted by smaller pie charts and subCSTs in blue font) but were manually re-classified based on their relative abundance profile that was not a good match to the other subCSTs (see Methods). CST IV-E contains a high to moderate relative abundance of *Prevotella* spp. with other species. Relative abundance data were used to plot the subCST IV-E pie chart (**Supplementary Table 5**).

Of the 95 study participants whose microbiomes did not match assigned subCSTs, iTaukei and other Pacific Islander ethnicities (68/74 (91.89%)) were significantly more likely to not match compared to Indo-Fijian and Other ethnicities (27/36 (75.00%); AOR: 3.78; 95% CI: 1.23-11.64; *P* = 0.0205) (**Supplementary Table 4**). Age group was not associated with a lack of matching to the assigned subCSTs.

### Within host paired endocervical and vaginal bacterial species composition differs for women with and without *C. trachomatis* infection

The relative abundance of bacterial taxa in the paired vaginal and endocervical microbiomes based on *Ct* status is shown in **Figure 4**. For the *Ct* negative endocervical and vaginal pairs, 19 (35.84%) of 53 pairs had differential species relative abundance between the two anatomic sites (boxed in black, **Figure 4**). The remaining 33 (63.46%) pairs, had relatively similar species relative abundance between the two sites. The presence of *L. crispatus*–dominated microbiomes in five (15.15%) of the 33 similar pairs were unique to the *Ct* negative group (boxed in green, **Figure 4**). The majority of the endocervical and vaginal microbiomes in the *Ct* negative group were dominated by *G. vaginalis* (23/52; 44.23%), *L. iners* (16/52; 30.76%) and *Prevotella spp.* (4/52; 7.69%).

**Figure 4.**
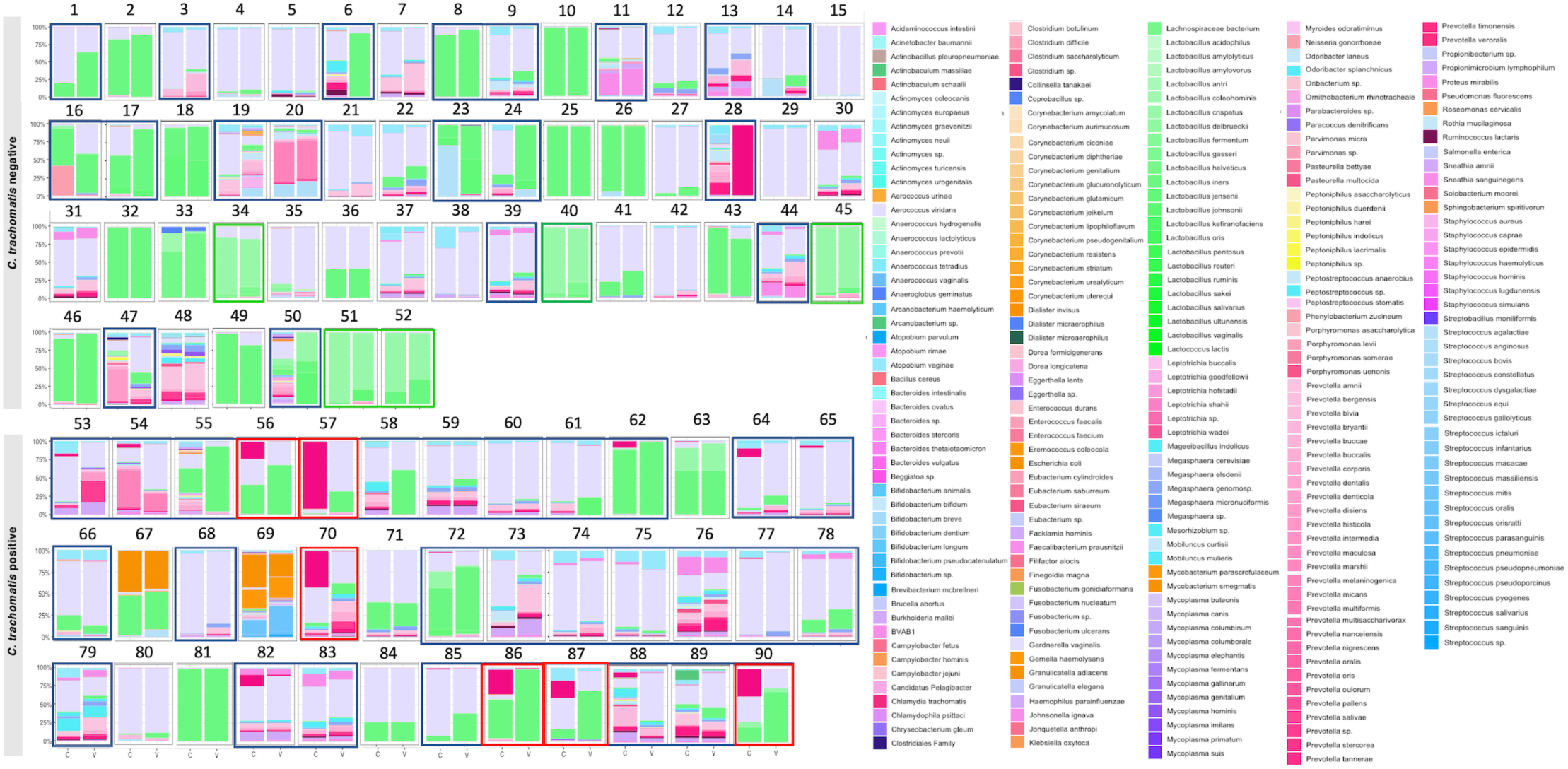
Relative abundance bar plots for *C. trachomatis* positive and for *C. trachomatis* negative paired endocervical and vaginal microbiomes. Only 90 pairs are shown due to lack of relative abundance data for two endocervical samples. Profiles in black boxes indicate paired endocervical and vaginal microbiomes with different microbial profiles. These microbiomes were dissimilar in species relative abundance for 19 (36.53%) of 52 *C. trachomatis* negative pairs and 31 (81.6%) of 38 positive pairs. Green boxes show *L. crispatus*-dominant paired endocervical and vaginal microbiomes with similar profiles, which were only found in *C. trachomatis* negative pairs (five (9.6%) of 52 pairs.) Red boxes show moderate to high relative abundance for *C. trachomatis* endocervical microbiomes with dissimilar profiles, which were only found in *C. trachomatis* positive pairs (six (15.79%) of 38 pairs.) C, endocervix; V, vagina.

The *Ct* positive paired endocervical and vaginal microbiomes were statistically more likely to have differential species relative abundance (31/38 (81.57%)) between the two sites compared to the *Ct* negative pairs (19/53 (35.84%); AOR: 7.93; 95% CI: 2.93-21.93; *P*<0.0001) (boxed in black, **Figure 4**). There was a higher relative abundance of *G. vaginalis* (18/38; 47.36%) compared to *L. iners* (7/38; 18.42%). Some profiles were also dominated by *Prevotella spp*. (3/38; 7.89%) and *E. coli* (2/38; 5.26%). Interestingly, in six (15.79%) of the 38 pairs (subjects 56, 57, 70, 86, 87 and 90), the endocervical microbiomes had a moderate to high relative abundance of *C. trachomatis* but not in their corresponding vaginal microbiomes.

### Higher microbial diversity and species evenness was found in the endocervix compared to the vagina

Alpha diversity was determined based on the relative abundance of species in a microbiome measured by Chao1 and Shannon index as shown in **Figure 5** using the Linear Decomposition Model (LDM)[32]. For vaginal microbiomes, iTaukei Pacific Islanders had a significantly higher alpha diversity compared to Indo-Fijian and other ethnicities in Fiji controlling for age and *Ct* (Chao1 *P*=0.0255, Shannon *P*=0.0322). Alpha diversity also varied significantly among the different subCSTs of the vaginal microbiome with higher diversity found among the subCST IV types (Chao1 *P*=2e-04, Shannon *P*=2e-04). Vaginal microbiomes with BV had a significantly higher diversity than those without BV controlling for *Ct* (Chao1 *P*=2e-04, Shannon *P*=0.0014), as did microbiomes with *Ct* comparted to those without *Ct* (Chao1 *P=*0.0072, Shannon *P*=0.0022) controlling for ethnicity and age.

**Figure 5.**
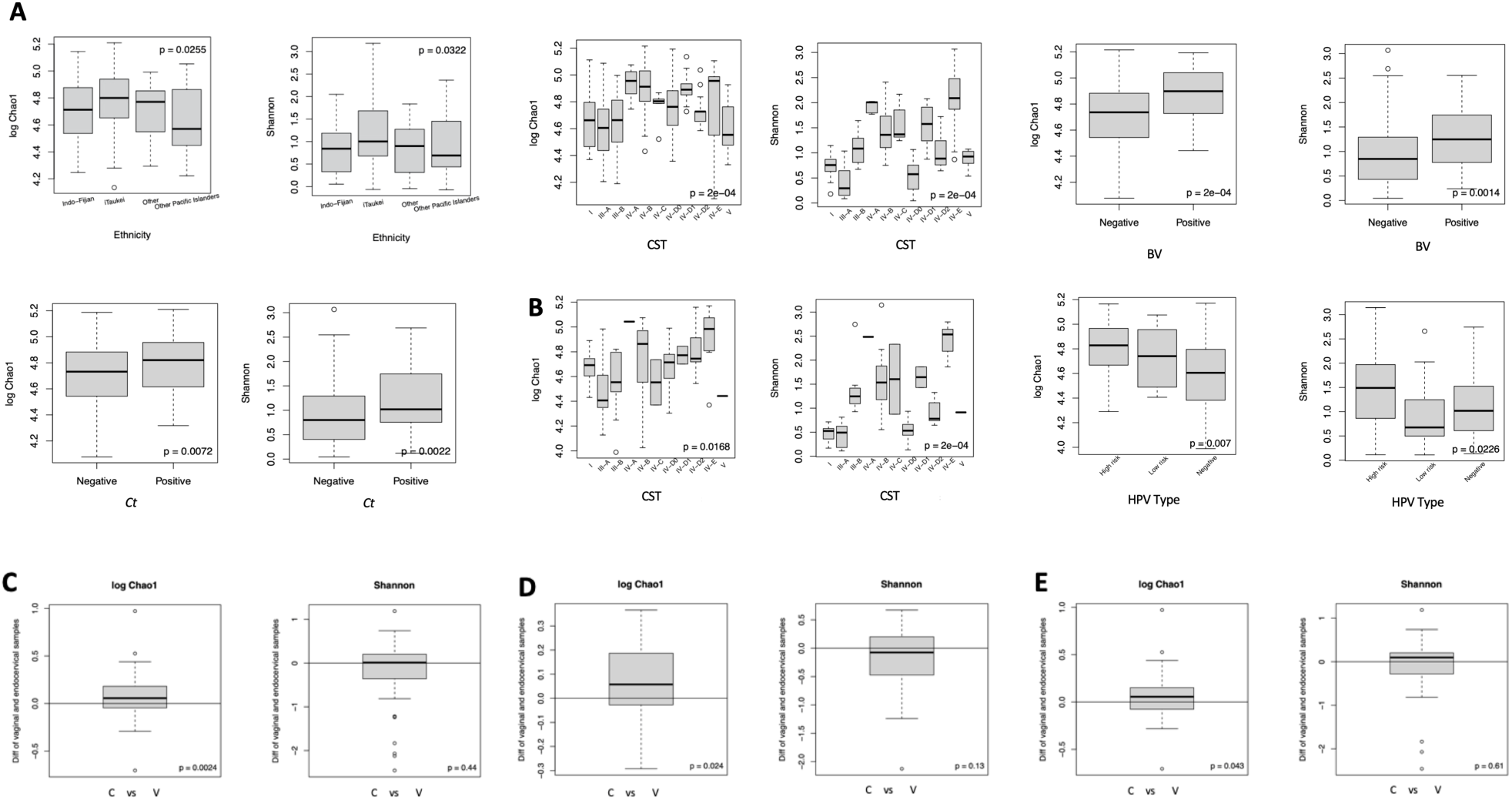
Alpha metrics for the vaginal and endocervical microbiomes with categorical variables and *P*-values estimated based on the Linear Decomposition Model (LDM; see Methods). **A**) Shown are the significant associations of the vaginal microbiomes with ethnicity, community state types (CST), including subCSTs, and bacterial vaginitis (BV). The significant association with *C. trachomatis (Ct)* status is shown. **B)** Significant associations with the endocervical microbiomes are community state types (CST) and high-risk human papilloma virus (hrHPV) types (see Supplementary Tables 1 and 2 for HPV risk types). **C**) Significant associations of alpha diversity metrics for paired endocervical and vaginal microbiomes are shown overall and independently for paired microbiomes based on *C. trachomatis* infection (**D**) and no *C. trachomatis* infection (**E**). Chao and Shannon indices were both used to measure alpha diversity (see Methods). C, endocervix; V, vagina.

Endocervical microbiomes had a significantly higher alpha diversity for all subCST IV classifications compared to other subCSTs (Chao *P*=0.0168, Shannon *P*=2e-04) (**Figure 5B)**. A significantly higher alpha diversity was also found for hrHPV types compared to lrHPV types and no HPV infection (Chao *P*=0.007, Shannon *P*=0.0226).

The comparison of alpha diversity between paired vaginal and endocervical microbiomes revealed that the two anatomic sites differed significantly based on Chao1 (*P*=0.0024) (**Figure 5C)** that was influenced by the presence of *Ct* (Chao1: *P*=0.024) (**Figure 5D)** and less so by the absence of *Ct* (Chao1: *P*=0.043) in the respective microbiomes (**Figure 5E).**

Beta diversity was employed to understand the divergence in microbial composition between samples measured by Bray-Curtis and the Jaccard metric. PCoA based on Bray-Curtis dissimilarities and Jaccard showed significant segregation of the vaginal microbiomes based on ethnic groups (*P*= 0.039 and *P*=0.0044), subCST categories (*P*= 2e-04 and *P*=2e-04), BV (*P*= 8e-04 and *P*=2e-04), Candida (*P*= 0.0068 and *P*=0.04. respectively) and *Ct* status (*P*= 0.012 and *P*=0.04, respectively) (**Figure 6)**. In contrast, the endocervical microbiomes were significantly segregated based on subCST IV categories (*P*= 2e-04 and *P*=2e-04, respectively) and *Ct* status (for Jaccard only*, P*=0.024) (**Figure 6)**.

**Figure 6.**
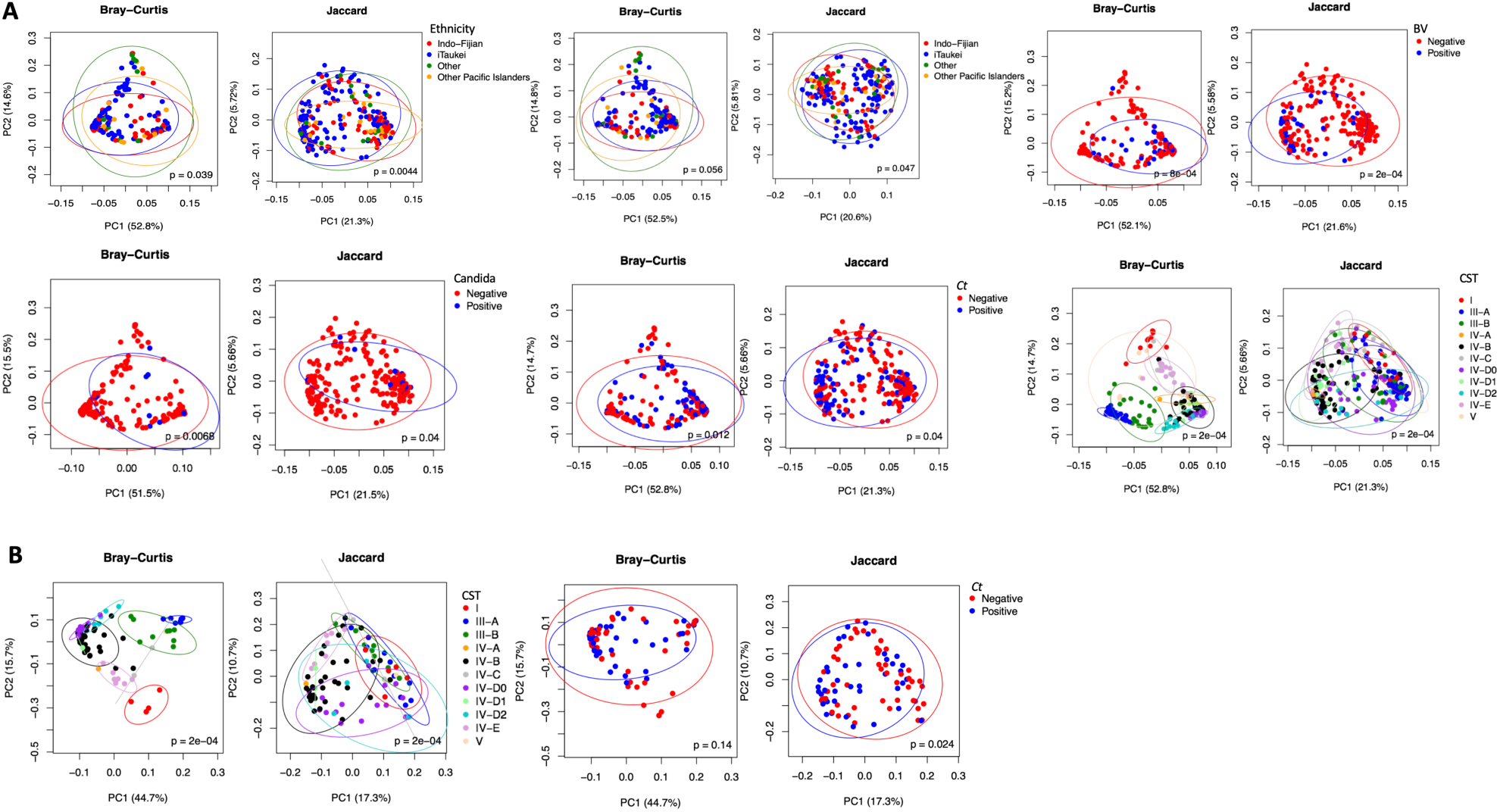
Beta-diversity metrics for the vaginal and endocervical microbiomes with categorical variables and *P*-values estimated based on Permutational Multivariate Analysis of Variance (PERMANOVA). A) Significant associations with the vaginal microbiomes are shown for ethnicity with and without controlling for *Ct* infection and age, community state types (CST), including subCSTs, bacterial vaginosis (BV), *Candida* and *C. trachomatis* (Ct) status. B) Significant associations are shown for community state types (CST), including subCSTs, and *C. trachomatis (Ct)* status with the endocervical microbiomes. Bray-Curtis and Jaccard indices were used to measure beta diversity (see Methods).

### Ethnicity, BV, *C. trachomatis* and anatomic site-specific associated species based on differential abundance and presence or absence of bacterial species

Based on differential abundance analysis using the Linear Decomposition Model (LDM) [32] vaginal microbiomes, iTaukei and Other Pacific Islander ethnicities had a significantly higher abundance of pathogenic bacteria including *S. amni, Leptotrichia goodfellowii,* and *Parvimonas sp.* compared to Indo-Fijian and Other ethnic groups in Fiji. Some species such as *D. microaerophilus, Prevotella sp.*, *Prevotella Salivae*, and *Bifidobacterium animalis* were associated with iTaukei ethnicity only (**Supplementary Figure 1)**. The high abundance of vaginal *L. crispatus* and *Actinomyces europaeus* were strongly associated with the Other ethnic group, which mainly comprised European, Chinese and mixed races. A different set of species were found to be enriched in the endocervical microbiomes, including *Ruminococcus lactaris*, *M indolicus*, *S. sanguinegens, Actinobaculum massiliae, Megashaera* sp. and *P. histicola,* and were significantly associated with iTaukei and/or Other Pacific Islander ethnicities. *L. aciophilus*, *B. dentium* and BVAB1 were associated with the other ethnicity group (**Supplementary Figure 2)**.

Based on LDM analysis, a total of 34 species were associated with BV, and all were significantly differentially abundant in BV positive compared to BV negative vaginal microbiomes (**Supplementary Figure 3)**. The majority of the species identified belonged to the genus *Dialister*, *Leptotrichia*, *Mageeibacillus*, *Megasphaera*, *Mobiluncus*, *Mycoplasma*, *Parvimonas*, *Prevotella*, *Porphyromonas* and *Sneathia*. Based on presence or absence, 33 species were significantly higher for microbiomes with BV, which largely overlapped with the differential abundance data with the exception of *L. crispatus* and *L gensenii,* which were significantly higher in BV negative microbiomes (**Supplementary Figure 3)**.

Differentially abundant species significantly associated with *Ct* positive vaginal microbiomes included *Prevotella sp*., *Tannerella forsythensis*, *L. goodfellowii*, *Lactobacillus pentosus*, *S. amnii*, *Megasphaera sp*. (**Supplementary Figure 4).** For presence-absence data*, L. crispatus* was significantly higher in presence in *Ct* negative biomes while *Sneathia sanguinegens*, *S. amnii*, *Tannerella forsythensis*, *Prevotella sp.*, *Lactobacillus pentosus*, *Megasphaera micronuciformis*, *Mageeibacillus indolicus*, *Prevotella pallens*, *Prevotella amnii* and *Bacteroides sp.* were significantly present in *Ct* positive microbiomes. No species were differentially abundant in the *Ct* infected endocervical microbiomes (data not shown.)

For *Ct* positive endocervical and vaginal paired microbiomes, 56 species were significantly differentially abundant in the endocervical compared to the paired vaginal microbiomes, and 12 species were differentially abundant in the vaginal compared to the endocervical biome (**Supplementary Figure 5**). Based on species presence or absence data, 71 species were significantly higher in the endocervical compared to vaginal biomes, and 19 species in the vaginal biomes compared to endocervical biomes (**Supplementary Figure 6**).

For *Ct* negative paired microbiomes, 26 species were significantly differentially abundant in the vaginal compared to endocervical microbiomes, and six species were differentially abundant in the endocervix (**Supplementary Figure 7**). For presence or absence data, 28 species were significantly higher in the vaginal biome compared to the endocervix, and four species in the endocervix compared to the vagina (**Supplementary Figure 8**).

When considering the anatomic site-specific differences regardless of *Ct* status, based on differential abundance, 67 species were significantly abundant in the endocervix compared to the paired vaginal microbiome, and 18 species were significantly abundant in the vagina compared to endocervix (**Supplementary Figure 9**). Based on presence or absence, 63 species were significantly higher in presence in the endocervix compared to the paired vaginal microbiomes, and 12 species in the vagina compared to the endocervix (**Supplementary Figure 10**).

### Paired endocervical and vaginal microbiomes show stable networks with some anatomic site variation depending on *C. trachomatis* status

To understand the bacterial community networks in the vaginal and endocervical microbiomes, we employed Pearson’s correlation coefficient with a cutoff of *P*<0.05 as statistically significant between different bacterial taxa in the respective microbiomes. Microbiota community clusters within the networks of endocervical and vaginal microbiomes were identified. Modality was calculated to measure the degree of stability within the densely connected communities in the network, which was found to be greater than 0.8 for all networks, suggesting a highly stable network of microbiomes [33]. Numbered groups represent a single species or related species (**Figure 7**); the species within the groups and clusters are shown in **Supplementary Table 6**. Nodes of the network, shown as enclosed groups, represent species where the node size denotes the abundance of those species in the community, imbuing a greater weight of interactions for a node. The edges represent mutualistic relationships within species of the community (**Figure 7**).

**Figure 7.**
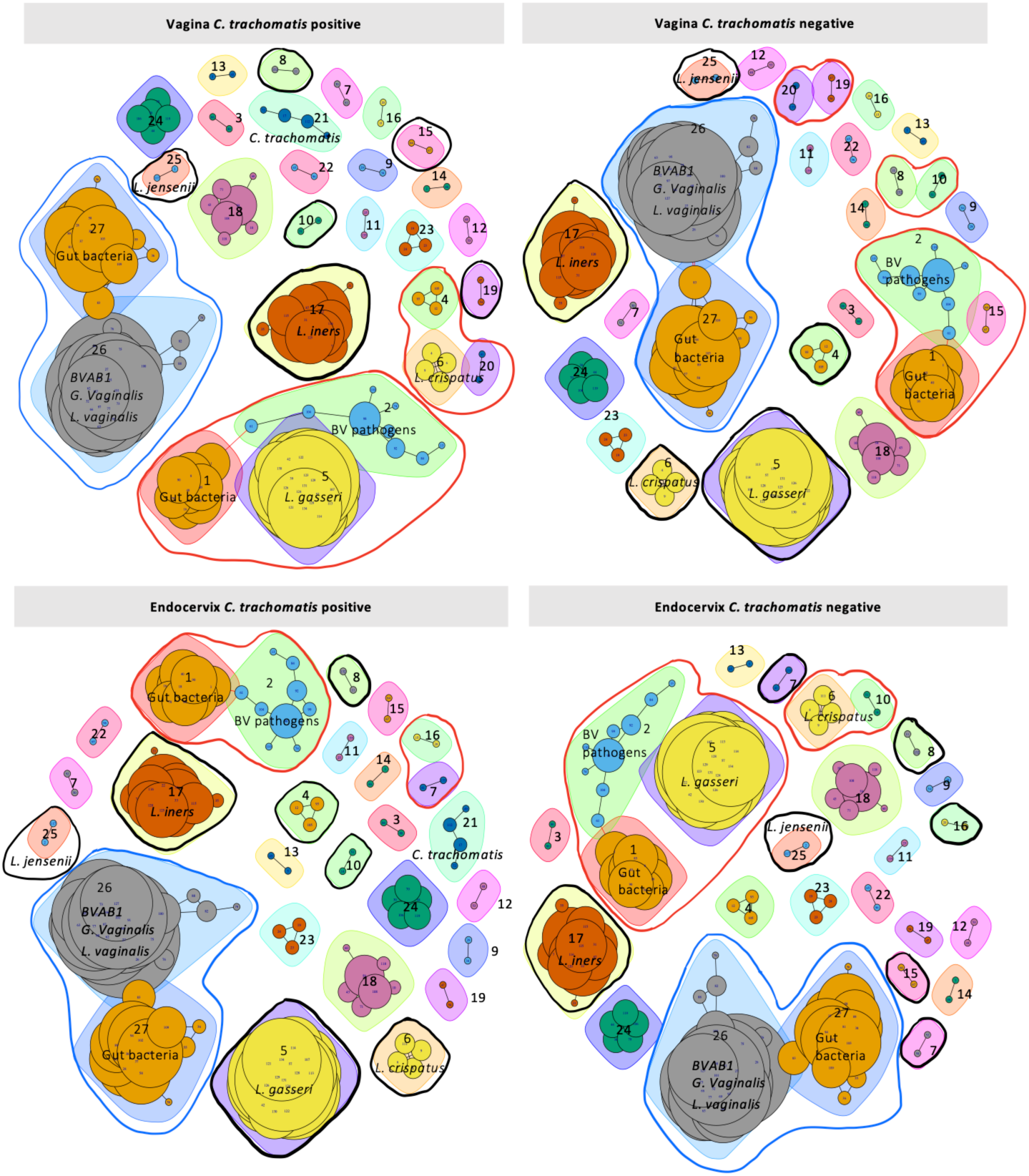
Network analysis based on Pearson’s correlation of the 92 *C. trachomatis* positive or negative paired endocervical and vaginal microbiomes. Graph partitioning was performed to identify microbial community clusters within each network, and “modularity” was calculated to measure the degree of stability within the densely connected compartments or communities (see Methods). Nodes represent two or more groups—single or related species shown as different colored shapes—where the larger the size of the node, the greater the weight of the interactions of the groups within the network. Red lines show nodes with differential groups between paired endocervical and vaginal *C. trachomatis* positive or negative networks and between endocervical and vaginal anatomic site networks. Black lines show differential independent groups between pairs and also between anatomic sites in the networks. Blue lines show similar nodes for all four *C. trachomatis* positive and negative endocervical and vaginal networks.

Nine clusters or nodes were identified among the *Ct* positive and negative paired endocervical and vaginal microbiomes (**Figure 7**). Cluster 1 consists of gut bacteria from group 27 genera (*Bacteroides*, *Bifidobacterium*, *Clostridium*, *Enterococcus*, *Fusobacterium*) and group 26 (BVAB1, *G. vaginalis*, *Lactobacillus vaginalis*, *Bacteroides*, *Corynebacterium*, *Dialister*, *Dorea*, *Enterococcus*, *Eubacterium* and *Fusobacterium*). The area for Cluster 1 was 35.60, 39.35, 36.94 and 37.04 for the *Ct* positive endocervix and vagina and *Ct* negative endocervix and vagina, respectively, where the larger the node, the more weight is given to the interaction in the network. Cluster 2 consisted of group 1 gut bacteria (*Anaerococcus*, *Bifidobacterium*, *Eggerthella* and *Faecalibacterium*) and group 2 BV pathogens (*Aerococcus*, *Corynebacterium*, *Dialister*, *E. coli*, *Finegoldia magna*, *Gemella*, *Klebsiella oxytoca*, *L. coleohominis*). Cluster 3 had groups 1 and 2 and group 5 (*L. gasseri*, *Leptotrichia*, *Bifidobacterium*, *Brevibacterium* and *Collinsella*). Cluster 4 comprised groups 1 and 2, and group 15 (*Corynebacterium* spp.). Cluster 5 is comprised of group 4 (*Actinomyces urogenitalis*, *Fusobacterium gonidiaformans*, and *Lachnospiraceae_bacterium*), group 6 (*L. crispatus* and *Actinomyces* spp.), and group 20 (*Bifidobacterium* and *Leptotrichia* spp.) Cluster 6, groups 2, 3 (*Enterococcus durans*, *Eubacterium saburreum*) and 27; Cluster 7, groups 19 (*Campylobacter jejuni* and *Clostridium difficile*) and 20; Cluster 8, groups 6 and 10 (*Aerococcus viridans* and *Chryseobacterium gleum*); and Cluster 9, groups 8 (*Leptotrichia shahii* and *Actinomyces turicensis*) and 10.

## Discussion

Major advances in our ability to analyze the microbial composition of vaginal microbiomes have revolutionized our understanding of the role they play in both health and disease. However, data on vulnerable populations from diverse ethnic and global populations are largely missing. Moreover, endocervical microbiomes and their relationship with the vaginal microbiome have barely been examined. The former is of particular importance because the endocervix but not the vagina is a primary site of infection for sexually transmitted *Ct*, *Ng* and HPV. In this study, we expanded on our previous pilot study [26] by evaluating 258 vaginal and 92 paired endocervical microbiomes for women of all ethnicities residing in Fiji. We characterized the microbial relative abundance, community composition and networks of these microbiomes and examined the unique relationship between paired endocervical and vaginal microbiomes.

Our 2020 study in Fiji showed that women of iTaukei ethnicity were significantly more likely to have *Ct* STIs than Indo-Fijians [3], and this was the case in the present study. Further, we found a significantly higher microbial diversity for iTaukei ethnicity compared to Indo-Fijian, Other Pacific Islanders and Other ethnicities. Many BV-associated bacteria such as *B. animalis*, *D. microaerophilus*, *Prevotella sp*., *P. salivae*, *S. amnii*, and *Parvimonas sp.* were also significantly associated with this ethnicity compared to other ethnicities. Further, there was a significant pattern of community composition for each ethnicity with iTaukei and Other Pacific Islanders having a diverse composition of anaerobes compared to a predominance of *Lactobacillus* spp. such as *L. crispatus* for Indo-Fijian and Other ethnicities. Such ethnicity- or race-based microbial clustering in microbiomes have also been reported for ethnic and racial subgroups of African descent and European/Caucasian background, respectively [21,22].

A number of Fijian women were diagnosed with BV and, as expected, there was a significant association with *Ct* infection. Vaginal microbiomes with BV also had a significantly higher diversity of microbial species and significant differences in community composition compared to microbiomes without BV as has been reported by [34,35]. Bacteria commonly associated with BV, including *Megaspheera* spp., *Sneathia* spp., *G. vaginalis* and *Prevotella* spp., were present at a significantly higher relative abundance compared to BV negative microbiomes. We also identified *A. intestenii*, *T. forsythensis* and *E. saburreum* as newly associated with BV. *A. intestenii* is an anaerobic intestinal commensal that has been found in the vaginal flora [36]. The other species are present in the oral biome and linked to periodontal disease and/or respiratory infections [37–39]. Their association with BV is not understood, although oral to genital transfer during sex is a possibility.

The Xpert CT/NG and wet prep tests on vaginal samples were able to detect *Ct*, *Ng*, *Tv* and *Candida*, while MSS detected each of these in addition to *Mg*. In our previous pilot study, *Mg* was found in 100% of our samples using the OneCodex database, although the sample size was small. In contrast, we detected only 3.5% and 7.6% in the vagina and endocervix, respectively, using VIRGO [24]. This discrepancy is not surprising because complete *Mg* genomes were used to build the VIRGO database and is therefore has higher specificity.

We were able to identify numerous HPV types using HPViewer [30], which provides much needed data on the distribution of genotypes among Pacific Islanders in the WPR. The only published study of HPV genotypes in PICTs identified hrHPV genotypes 16, 18, 33, 39, 45, and 52 among cervical cancer specimens along with lrHPV genotypes 54 and 67 that are infrequently found in these malignancies [40,41]; no data were available for women without cervical cancer. hrHPV 16, 18, 39 and 52 were also found in our Fijian population. Of the common lrHPV types present in the population, HPV 57 has been commonly associated with oral papillomas [42]. A possible oral to genital auto-inoculation or sexual transmission could explain the unexpected presence of this genotype in genital microbiomes.

Although no significant association of endocervical hrHPV genotypes with ethnicity, age or *Ct* infection was found, not surprisingly, these high-risk types were associated with a higher microbial diversity compared to low-risk types or no infection in the endocervical microbiome. This finding is consistent with two recent studies in China that showed a significantly increased alpha diversity in the cervix of women with HPV compared to those without infection [43,44].

Many endocervical (41.3%) and vaginal (36.04%) microbial relative abundance profiles did not match their assigned subCST based on the 13 VALENCIA reference centroids. By manually comparing the species relative abundance of each microbiome with the assigned subCSTs, we were able to develop four new subCSTs for CST IV—namely IV-D0, IV-D1, IV-D2 and IV-E—that provide a more specific classification for Fijian microbiomes. Many of the microbiomes classified as IV-B required reclassification as IV-D0, IV-D1 and IV-D2, and microbiomes originally assigned as CST III-B, IV-A, IV-B, IV-C0, IV-C1 or IV-C4 were re-classified as CST IV-E. The need to expand CST classifications is supported by a 2022 study of ethnic Han Chinese where a predominance of *L. helveticus* was discovered based on 16S rRNA sequences of vaginal microbiomes not reflected in VALENCIA reference centroids [31]. They proposed a new CST VI to represent communities that are dominated by this *Lactobacillus* sp. Additional subCSTs and CSTs will likely be required to further define endocervical and vaginal microbial landscapes as more diverse and globally represented populations are studied.

The new subCSTs IV-D0, IV-D1, IV-D2 and IV-E had a predominance of *G. vaginalis, L. iners* and *Prevotella* spp. along with other BV-associated pathogens. Many of these microbiomes may have predisposed to sexually transmitted *Ct* and/or HPV infections in this population. However, while there was no significant association between new subCSTs that did not match assigned subCSTs and *Ct* infection, there was a significant association of unmatched subCSTs with iTaukei and Other Pacific Islander ethnicities compared to Indo-Fijian and Other ethnicities.

The paired endocervical and vaginal microbiomes had unique attributes depending on *Ct* status. Among *Ct* negative pairs, while a little over a third had differential species relative abundance between the two anatomic sites, 15.15% of the pairs with similar species were dominated by *L. crispatus* with a slightly greater relative abundance in the endocervix. This is in contrast to a study where uninfected Chinese women were found to have a higher abundance of this species in the vagina compared to the endocervix [45]. Multiple investigations have shown that *L. crispatus* produces D-lactic acid and bacteriocins that may trap or inhibit pathogen attachment to host cells, protecting against STIs and BV to maintain microbiome eubiosis [46–48]. Studies of South African adolescents and Italian women also found that *L. crispatus*-dominated microbiomes were less likely to be associated with *Ct* infection [13,49].

Among *Ct* positive pairs, over 80% had differential species relative abundance with 72 species showing a significant difference between the endocervical and vaginal microbiomes. Not surprisingly, *Ct* positive pairs were significantly more likely to have differential species relative abundance between the two anatomic sites compared to *Ct* negative pairs. Further, the *Ct* infected endocervix had a higher microbial diversity compared to the vagina. These data are consistent with our previous pilot study [26] and a study of South African adolescents that also examined endocervical and vaginal microbiomes [13]. A recent study of HPV infections found similar results [50]. In addition, the high dominance of *G. vaginalis* along with other BV-associated pathogens such as *Prevotella*, *Megasphaera* and *Atopobium* among *Ct* positive microbiomes in our study has also been reported in South Africa and Italy, although 16Ss rRNA-based sequencing resolution was used in those studies [13,49]. The relatively lower dominance of *L. iners* (18.4%) compared to the *Ct* negative pairs (30.7%) could be explained by the fact that *L. iners* lacks the essential amino acid synthesis repertoire, forcing it to rely on exogenous host amino acids, which would make it more sensitive to the community composition and perhaps less successful in competing with other microbes [51]. However, *L. iners* has been shown in other studies to be associated with *Ct* infection and is more prevalent among women of African descent than other racial or ethnic groups [52–54].

Six (15.79%) of the *Ct* positive endocervical microbiomes were dominated by a moderate to high relative abundance of *Ct,* but not in the paired vaginal microbiomes. Although the columnar epithelial cells of the endocervix are the target for *Ct* infection, this level of abundance has not previously been reported, and was a unique occurrence in Pacific Islanders. Indeed, the few studies that evaluated the endocervical microbiome did not find this level of *Ct* abundance among, for example, Malaysian, Chinese, Indian, Italian and South African women [13,49,55]. A larger prospective study is required to assess the frequency of this finding in Pacific Islander ethnicities and the significance in terms of microbial composition and disease pathogenesis.

Nine clusters or nodes were identified in the paired endocervical and vaginal microbiome networks with shifts in size and composition depending on *Ct* status and anatomic site. However, Cluster 1, comprising a combination of gut bacteria and BV-associated species, was present in all four networks. This finding reflects the very nature of the Pacific Islander microbiomes in that they tend to be highly diverse and rich in pathogenic bacteria regardless of *Ct* status, which is consistent with our previous findings [26]. However, this cluster was appreciably smaller in the *Ct* positive endocervical network compared to all others and had no clusters that included *Lactobacillus* spp. (i.e., groups 5, 6, 17 or 25) while two of the three clusters in the *Ct* positive vaginal microbiome contained *L crispatus* and *L. gasseri*, respectively. Since *Ct* infects the endocervical epithelium but, in the vagina, *Ct* organisms are either inside sloughed cells or loose, the influence of *Ct* on each anatomic site might be distinct. Nodes containing *Lactobacillus* spp. may interact with BV-associated species and gut bacteria to dampen the effects of these pathogenic species or vice versa. Further, the high abundance of *G. vaginalis* may form biofilms but also interact with other BV-associated spp. to generate polymicrobial biofilms that increase the risk and persistence of *Ct* and other STIs [56,57]. *Ct* infected endocervical cells may actually work to dissociate or prevent *Lactobacillus* spp. interactions. This would be consistent with the smaller nodes that lack *Lactobacillus* spp. in the *Ct* positive endocervical networks compared to the *Ct* positive vaginal networks. Further, this is consistent with our findings that the *Ct* negative endocervical network was similar to the *Ct* positive vaginal network. These network interactions are therefore likely driven by the microbial milieu of gut and BV-associated bacteria that are influenced by *Ct* infection in the endocervix and not by the simple presence of *Ct*, free or otherwise, as in the vagina. Additional studies will be needed to tease out these pathogenic interactions.

## Conclusions

*Ct* is the most common sexually transmitted bacterium in the world today with a steady increase in cases since the 1990s. Endocervical microbiomes and their relationship with the vaginal microbiome have rarely been examined in the context of *Ct*. Further, our understanding of these microbiomes among vulnerable groups from diverse ethnic and global populations are lacking. Here, 258 vaginal and 92 paired endocervical samples from women of diverse ethnicities in Fiji were evaluated using MSS. We found that Pacific Islander ethnicities and women under 25 years of age were significantly associated with *Ct* infection. Over 37% of the microbiomes could not be classified into subCSTs using the VALENCIA reference centroids. We therefore developed four new subCSTs IV-D0, IV-D1, IV-D2, and IV-E—variably dominated by *G. vaginalis, Prevotella* spp. and *L. iners*—to better classify these microbiomes. Not surprisingly, *Ct* positive endocervical and paired vaginal microbiomes were significantly more likely to have differential species relative abundance than negative pairs. Further, the endocervix had a higher alpha diversity, which was also associated with hrHPV genotypes, compared to paired vaginal microbiome. Despite the presence of gut and BV-associated bacterial clusters in both *Ct* positive and negative paired microbiome networks, there was a unique microbial network in the *Ct* positive endocervix with smaller bacterial nodes and no interactions with *Lactobacillus* spp., suggesting a distinct influence on this microbiome that might be explained in part by *Ct* infection of the endocervical columnar epithelium and *G. vaginalis* generated polymicrobial biofilms that provoke pathogenic disease in the host tissue. Our findings expand on the existing repertoire of endocervical and vaginal microbiomes, filling in the gaps in our knowledge about Pacific Islanders. However, additional research is needed to evaluate network interactions, especially for biofilms in the presence or absence of *Ct* infection, and prospective studies to further define causal associations that could lead to successful interventional strategies.

## Methods

### Sample collection and patient characteristics

Women attending the Fijian Ministry of Health and Medical Services (MoHMS) health centers and outreach clinics in the Central Division, Viti Levu, Fiji, were enrolled after informed consent as part of a parent study as previously described [3]. Human subject’s approval was obtained from the Institutional Review Boards of the Fiji Ministry of Health and Medical Services and the University of California San Francisco. Briefly, women of various ethnicities residing in Fiji aged 18 to 40 years were enrolled unless they had HIV, untreated syphilis, a diagnosis of cancer or had received antibiotics within the prior month. The parent study provided de-identified vaginal *Ct* positive and negative samples for a total of 258; 92 samples were available from both vaginal and endocervical sites of the same women. All swabs were collected by a trained physician and stored in SWAB/A-50 buffer (Cepheid, Sunnyvale, CA) and/or 1 mL of M4 media (Copan. Murrieta, CA). Vaginal samples were tested using the Xpert CT/NG assay (Cepheid), and endocervical samples were screened for *Ct* using an in-house qPCR as we described [26]. Samples were additionally tested for *Tv* and *Candida* by wet prep. A diagnosis of bacterial BV was made using three or more Amsel criteria [29]: homogeneous vaginal discharge, >20% clue cells on wet prep, vaginal pH ≥4.5, and fishy amine odor when KOH was applied to vaginal material on a glass slide.

### Sample processing, metagenomic shotgun sequencing and taxonomic analysis

Genomic (g)DNA was extracted from endocervical and vaginal swab samples, enriching for microbial DNA using a lysozyme cocktail method prior to gDNA purification using the QIAmp DNA Mini Kit (Qiagen, Germantown, MD) as we described [26]. DNA concentrations were measured by Qubit^TM^ dsDNA BR assay kit (Invitrogen, Carlsbad, CA). Metagenomic shotgun sequencing libraries were constructed using Illumina Nextera XT kits and sequenced using 150 nucleotide paired-end reads on an Illumina Novaseq PE150 platform.

Raw sequence data was processed using BBDuk v38.96 for Illumina adapter and quality trimming [58]. Read regions were trimmed to an average quality score of >10, and read ends were excised if their quality scores were <10. Human read contaminants were removed using BBSplit v38.96 [58]. Version 38 of the human genome (RefSeq: GCF_000001405.26) was used to build a reference for contaminant removal. Ambiguous reads or reads that were not clearly human in origin were preserved for downstream processing. Contaminant free read data were then profiled using Metaphlan v3.0.14 for species identification and Humann v3.0.1 for pathway and functional identification [59,60]. VIRGO [24] was used on the post-processed vaginal and endocervical microbial reads to determine the taxonomic composition, functional pathways and protein attributes. The number of confirmed reads for *Mg* obtained from VIRGO were divided by the total number of microbial reads for that sample, excluding human reads, and then multiplied by 1 million to arrive at a value for reads per million (RPM) for that pathogen [61]. HPViewer [30] was used to identify and quantify the presence of HPV types in the vaginal and endocervical metagenomes.

### Community state type (CST) assignments

The abundance data generated from VIRGO [24] was used for classification of the vaginal and endocervical microbiomes into community state types (CSTs) using a nearest-centroid based algorithm called VALENCIA [25]. CSTs were assigned based on the highest similarity score of a microbial profile for the 15 reference centroids, ranging from 0.0 (no shared taxa) to 1.0 (all taxa shared at the same relative abundance).

Based on our previous study [26] and a significantly larger sample size in the present study, there was a high discrepancy between the species composition of the pacific islander microbiomes and the subCST scores assigned by VALENCIA. We therefore manually compared the VALENCIA assigned subCSTs with the species composition of the 258 vaginal and 92 endocervical microbiomes to determine whether any new CST subtypes could be developed based on the existing five CSTs developed by Ravel et al. [15] and the 13 reference subCSTs included in VALENCIA.

### Statistical analysis and visualization

Bivariate and multivariate analyses of the association between covariates and *C. trachomatis* were performed using logistic regression models controlling for confounding variables. Results were expressed as odds ratios and adjusted odds ratios with 95% CI. All analyses were performed using R version 4.0.3 including the stats (4.0.3), dplyr (1.0.8) and broom (1.0.1) packages [62]. A significance level of <0.05 was used for all statistical analyses.

Microbiome analysis was performed using R software packages [62] Vegan (v.1.4-5) [63], Adespatial (v.0.3-14) [64], ggplot2 (v.3.3.5) [65], CGPfunctions (v.0.6.3) [66], FSA (v.0.9.3) [67], and ggpubr (v.0.4.0)[68] in addition to LDM [32]. Differences in Shannon indices between paired endocervical and vaginal microbiomes were estimated using odds ratio or risk ratios with 95% CI and tested by the paired two-sample T test, chi-square test of association, or Fishers exact test when cells were small. Differences in Shannon indices between *Ct* positive and negative microbiomes were tested by the unpaired two-sample T test. The relative abundance of the top 25 species within each patient’s vaginal and endocervical microbiome were visualized using the ggplot2 (v.3.3.5) library [65]. Principal coordinate analysis was used to visualize the clustering of microbiome samples based on the Bray-Curtis distances, with statistical hypothesis testing performed by Permutational Multivariate Analysis of Variance (PERMANOVA) (see below). Statistical comparisons across the *Ct* positive and negative groups was performed using Kruskal-Wallis test and LDM.

Correlations between different bacterial taxa in the vaginal and endocervical microbiomes based on *Ct* status were determined by computing Pearson’s correlation coefficients (with a *P-*value cut off of 0.05) using R packages igraph (v. 1.3.0) [33], Hmisc (v. 4.6-0) [69] and Matrix (v. 1.4-1) [70]. Using these packages, graph partitioning of the networks was done to identify community clusters within each network, and ‘modularity’ was calculated to measure the degree of stability within the densely connected communities.

The differential relative abundance association testing was primarily carried out by the LDM method controlling for confounding variables [32,71]. It is based on linear models of regressing individual taxon data on the covariate that we wish to test. The taxon data are taken to be relative abundances, arcsin-root transformed relative abundances, and presence-absence statuses, and the results from analyzing all these data scales are combined to produce overall tests of differential abundance at both the species and community levels. LDM bases the inference on a permutation procedure, which is important to preserve the data structure and thus data correlations in our data. The permutation procedure was performed by stratifying the groups of subjects who each had the same number of visits. For each group of subjects, the covariate values from a subject were shuffled as a set with the covariate values from another subject. In addition, the covariate values within a subject were also shuffled. The species-level analysis detected associated species at the nominal FDR level of 10%. The community-level analysis aggregated information from the species-level analysis. Both the global *P*-value and individual *P*-values in the boxplots were produced by the LDM. The LDM for presence-absence analysis [71] is based on rarefaction, and the LDM aggregates information from the all rarefied taxa count table. The *P*-values for testing the association of each beta diversity metric with a variable was based the PERMANOVA method [32,72] using the same permutation procedure described above. *P*-values were adjusted for multiple testing (testing multiple species) by the Sandve’s algorithm, which is integrated in the LDM package [71].

### Data availability

The metagenomic shotgun sequences as human read-removed data are available as FASTQ files along with associated metadata in the National Center for Biotechnology Information (NCBI) Sequence Read Archive (SRA) under the BioProject ID PRJNA982400 (https://www.ncbi.nlm.nih.gov/bioproject/?term=PRJNA982400). The Supplementary Tables provide data per unique ID# on age, ethnicity, STI infections, Candida, BV, and HPV genotypes; metagenomic shotgun sequencing results and quality control statistics for each sample; relative abundance for the top 28 species and by genera for each microbiome; average of the relative abundance data for each species of the newly assigned sub community state types (subCSTs) for microbiomes meeting the criteria for the new subCSTs; and microbial composition for anatomic networks. **Supplementary Table 7** provides the accession numbers for all metagenomic samples in NCBI. All scripts for the associated figures can be found at https://github.com/ddeanlab/Microbial-PI-Ct-MSS.

## Acknowledgements

We thank the parent study for providing the deidentified data for this study and Fijian colleagues: Dr. Rachel Devi; Dr. Kinisimere Nadredre; Dr. Mere Kurulo; and Dr. Darshika Balak. This research was supported by Public Health Service grant from the National Institutes of Health R01 AI151075 (to DD and TDR).

## Supplementary Information

### Supplementary Figures

**Supplementary Figure 1.** Species significantly associated with ethnicity based on differential abundance (A) or their presence or absence (B) for the vaginal microbiomes. Linear Decomposition Model (LDM) was used for statistical associations (see Methods).

**Supplementary Figure 2.** Species significantly associated with ethnicity based on differential abundance (A) or their presence or absence (B) for the endocervical microbiomes. Linear Decomposition Model (LDM) was used for statistical associations (see Methods).

**Supplementary Figure 3.** Species significantly associated with bacterial vaginosis (BV) for the vaginal microbiome based on differential abundance (A) or their presence or absence (B). Linear Decomposition Model (LDM) was used for statistical associations (see Methods). Note that BV cannot be measured for the endocervix.

**Supplementary Figure 4.** Species significantly associated with *C. trachomatis* compared to no infection for the vaginal microbiomes based on differential abundance (A) or their presence or absence (B). Linear Decomposition Model (LDM) was used for statistical associations (see Methods). No *C. trachomatis* associated species were identified for the endocervix.

**Supplementary Figure 5.** Species significantly associated with anatomic site based on differential abundance for the paired *C. trachomatis* infected endocervical and vaginal microbiomes. Linear Decomposition Model (LDM) was used for statistical associations (see Methods). C, endocervix; V, vagina.

**Supplementary Figure 6.** Species significantly associated with anatomic site based on presence or absence for the paired *C. trachomatis* infected endocervical and vaginal microbiomes. Linear Decomposition Model (LDM) was used for statistical associations (see Methods). C, endocervix; V, vagina.

**Supplementary Figure 7.** Species significantly associated with anatomic site based on differential abundance for the paired *C. trachomatis* uninfected endocervical and vaginal microbiomes. Linear Decomposition Model (LDM) was used for statistical associations (see Methods). C, endocervix; V, vagina.

**Supplementary Figure 8.** Species significantly associated with anatomic site based on presence or absence for the paired *C. trachomatis* uninfected endocervical and vaginal microbiomes. Linear Decomposition Model (LDM) was used for statistical associations (see Methods). C, endocervix; V, vagina.

**Supplementary Figure 9.** Species significantly associated with anatomic site based on differential abundance for the paired endocervical and vaginal microbiomes. Linear Decomposition Model (LDM) was used for statistical associations (see Methods). C, endocervix; V, vagina.

**Supplementary Figure 10.** Species significantly associated with anatomic site based on presence or absence for the paired endocervical and vaginal microbiomes. Linear Decomposition Model (LDM) was used for statistical associations (see Methods). C, endocervix; V, vagina.

### Supplementary Tables

**Supplementary Table 1.** Participant chararchteristics including infection with various sexually transmitted infections, bacterial vaginosis (BV) and *Candida*.

**Supplementary Table 2.** Metagenomic shotgun sequencing (MSS) data for *Neisseria gonorrhoeae* and *Mycoplasma genitalium* confirmed by VIRGO; *Trichomonas vaginalis* and *Candida albicans* confirmed by MetaPhlAn v3.0; and HPV by HPViewer.

**Supplementary Table 3.** Metagenomic shotgun sequencing results and quality control statistics.

**Supplementary Table 5.** Average of the relative abundance data for each species of the newly assigned sub community state types (subCSTs).

**Supplementary Table 4.** New classification of subCSTs for Pacific Islanders, and metrics for age, ethnicity and C. trachomatis infection status.

**Supplementary Table 6.** List of all species within the groups and clusters of vaginal and endocervical microbiome networks (see Figure 7).

**Supplementary Table 7.** Accession numbers for all metagenomic samples submitted to NCB

## References

1. Global health sector strategy on Sexually Transmitted Infections, 2016-2021 [Internet]. World Health Organization. 2016. https://www.who.int/publications/i/item/WHO-RHR-16.09. Accessed 2 Apr 2023.

2. Programmes STI. Global progress report on HIV, viral hepatitis and sexually transmitted infections, 2021 [Internet]. World Health Organization. 2021. http://www.who.int/publications/i/item/9789240027077. Accessed 2 Apr 2023.

3. Svigals V, Blair A, Muller S, Khan AS, Faktaufon D, Kama M, et al. Hyperendemic *Chlamydia trachomatis* sexually transmitted infections among females represent a high burden of asymptomatic disease and health disparity among Pacific Islanders in Fiji. PLOS Neglected Tropical Diseases. 2020. p. e0008022. doi:10.1371/journal.pntd.0008022.

4. Cliffe SJ, Tabrizi S, Sullivan EA, Pacific Islands Second Generation HIV Surveillance Group. Chlamydia in the Pacific region, the silent epidemic. Sex Transm Dis. 2008;35:801–6.

5. van Gemert C, Stoove M, Kwarteng T, Bulu S, Bergeri I, Wanyeki I, et al. Chlamydia prevalence and associated behaviours among female sex workers in Vanuatu: results from an integrated bio-behavioural survey, 2011. AIDS Behav. 2014;18:2040–9.

6. Vallely LM, Toliman P, Ryan C, Rai G, Wapling J, Tomado C, et al. Prevalence and risk factors of *Chlamydia trachomatis*, *Neisseria gonorrhoeae*, *Trichomonas vaginalis* and other sexually transmissible infections among women attending antenatal clinics in three provinces in Papua New Guinea: a cross-sectional survey. Sex Health. 2016;13:420–7.

7. Greub G. Chlamydial Infection: A Clinical and Public Health Perspective. Emerg Infect Dis. 2014;20:1266. doi: 10.3201/eid2007.140490.

8. Mania-Pramanik J, Kerkar S, Sonawane S, Mehta P, Salvi V. Current *Chlamydia trachomatis* Infection, A Major Cause of Infertility. J Reprod Infertil. 2012;13:204–10.

9. Peterman TA, Newman DR, Maddox L, Schmitt K, Shiver S. Risk for HIV following a diagnosis of syphilis, gonorrhoea or chlamydia: 328,456 women in Florida, 2000– 2011. Int J STD AIDS. SAGE Publications; 2015;26:113–9.

10. Koskela P, Anttila T, Bjørge T, Brunsvig A, Dillner J, Hakama M, et al. *Chlamydia trachomatis* infection as a risk factor for invasive cervical cancer. Int J Cancer. 2000;85:35–9.

11. Sorchik R, Than J, Carter K, Linhart C, Haberkorn G. Fertility trends in Pacific Island countries and territories. Statistics for Development Division, Pacific Community. 2019. https://sdd.spc.int/digital_library/fertility-trends-pacific-island-countries-and-territories2019. Accessed 2 Apr 2023.

12. van der Veer C, Bruisten SM, van der Helm JJ, de Vries HJC, van Houdt R. 2017. The Cervicovaginal Microbiota in Women Notified for *Chlamydia trachomatis* Infection: A Case-Control Study at the Sexually Transmitted Infection Outpatient Clinic in Amsterdam, The Netherlands. Clin infect Dis. 2017;64:24–31.

13. Balle C, Lennard K, Dabee S, Barnabas SL, Jaumdally SZ, Gasper MA, et al. Endocervical and vaginal microbiota in South African adolescents with asymptomatic *Chlamydia trachomatis* infection. Sci Rep. 2018;8:11109. doi:10.1038/s41598-018-29320-x

14. Brotman RM. Vaginal microbiome and sexually transmitted infections: an epidemiologic perspective. J Clin Invest. 2011;121:4610–7.

15. Ravel J, Gajer P, Abdo Z, Schneider GM, Koenig SSK, McCulle SL, et al. Vaginal microbiome of reproductive-age women. Proc Natl Acad Sci U S A. 2011;108 Suppl 1:4680–7.

16. Di Paola M, Sani C, Clemente AM, Iossa A, Perissi E, Castronovo G, et al. Characterization of cervico-vaginal microbiota in women developing persistent high-risk Human Papillomavirus infection. Sci Rep. 2017;7:10200. doi:10.1038/s41598-017-09842-6.

17. Brotman RM, Shardell MD, Gajer P, Tracy JK, Zenilman JM, Ravel J, et al. Interplay between the temporal dynamics of the vaginal microbiota and human papillomavirus detection. J Infect Dis. 2014;210:1723–33.

18. Tamarelle J, Thiébaut ACM, de Barbeyrac B, Bébéar C, Ravel J, Delarocque-Astagneau E. The vaginal microbiota and its association with human papillomavirus, *Chlamydia trachomatis*, *Neisseria gonorrhoeae* and *Mycoplasma genitalium* infections: a systematic review and meta-analysis. Clin Microbiol Infect. 2019;25:35– 47.

19. Zhou X, Brown CJ, Abdo Z, Davis CC, Hansmann MA, Joyce P, et al. Differences in the composition of vaginal microbial communities found in healthy Caucasian and black women. ISME J. 2007;1:121–33.

20. Gupta VK, Paul S, Dutta C. Geography, Ethnicity or Subsistence-Specific Variations in Human Microbiome Composition and Diversity. Front Microbiol. 2017;8:1162. doi:10.3389/fmicb.2017.01162.

21. Borgdorff H, van der Veer C, van Houdt R, Alberts CJ, de Vries HJ, Bruisten SM, et al. The association between ethnicity and vaginal microbiota composition in Amsterdam, the Netherlands. PLoS One. 2017;12:e0181135. DOI: 10.1371/journal.pone.0181135.

22. Fettweis JM, Brooks JP, Serrano MG, Sheth NU, Girerd PH, Edwards DJ, et al. Differences in vaginal microbiome in African American women versus women of European ancestry. Microbiology. 2014;160:2272–82.

23. MacIntyre DA, Chandiramani M, Lee YS, Kindinger L, Smith A, Angelopoulos N, et al. The vaginal microbiome during pregnancy and the postpartum period in a European population. Sci Rep. 2015;5:8988. doi.org/10.1038/srep08988.

24. Ma B, France MT, Crabtree J, Holm JB, Humphrys MS, Brotman RM, et al. A comprehensive non-redundant gene catalog reveals extensive within-community intraspecies diversity in the human vagina. Nat Commun. 2020;11:940. doi: 10.1038/s41467-020-14677-3.

25. France MT, Ma B, Gajer P, Brown S, Humphrys MS, Holm JB, et al. VALENCIA: a nearest centroid classification method for vaginal microbial communities based on composition. Microbiome. 2020;8:166. doi:10.1186/s40168-020-00934-6.

26. Bommana S, Richards G, Kama M, Kodimerla R, Jijakli K, Read TD, et al. Metagenomic Shotgun Sequencing of Endocervical, Vaginal, and Rectal Samples among Fijian Women with and without *Chlamydia trachomatis* Reveals Disparate Microbial Populations and Function across Anatomic Sites: a Pilot Study. Microbiol Spectr. 2022;10:e0010522. doi: 10.1128/spectrum.00105-22.

27. McKinney IM. Statistical News. Encyclopedia of Statistical Sciences. Hoboken, NJ, USA: John Wiley & Sons, Inc. 2006. https://onlinelibrary.wiley.com/doi/10.1002/0471667196.ess2539.pub2. Accessed 2 Apr 2023.

28. Naidu V, Matadradra A, Sahib M, Osborne J. Fiji: The challenges and opportunities of diversity. The University of the South Pacific. 2013. http://repository.usp.ac.fj/7723/1/Fiji_the_Challenges_and_Opportunities_of_Diversity_.pdf.

29. Amsel R, Totten PA, Spiegel CA, Chen KC, Eschenbach D, Holmes KK. Nonspecific vaginitis. Diagnostic criteria and microbial and epidemiologic associations. Am J Med. 1983;74:14–22.

30. Hao Y, Yang L, Galvao Neto A, Amin MR, Kelly D, Brown SM, et al. HPViewer: sensitive and specific genotyping of human papillomavirus in metagenomic DNA. Bioinformatics. 2018;34:1986–95.

31. Li H, Jiang J, Nie C, Xiao B, Li Q, Yu J. Community structure and ecological network’s changes of vaginal microbiome in women right after delivery. Front Pediatr. 2022;10:750860. doi: 10.3389/fped.2022.750860.

32. Hu Y-J, Satten GA. Testing hypotheses about the microbiome using the linear decomposition model (LDM). Bioinformatics. 2020;36:4106–15.

33. Csardi G, Nepusz T, Others. The igraph software package for complex network research. InterJournal, complex systems. 2006;1695:1–9.

34. Srinivasan S, Hoffman NG, Morgan MT, Matsen FA, Fiedler TL, Hall RW, et al. Bacterial communities in women with bacterial vaginosis: high resolution phylogenetic analyses reveal relationships of microbiota to clinical criteria. PLoS One. 2012; 7:e37818. doi: 10.1371/journal.pone.0037818.

35. Onderdonk AB, Delaney ML, Fichorova RN. The human microbiome during bacterial vaginosis. Clin Microbiol Rev. 2016;29:223–38.

36. D’Auria G, Galán J-C, Rodríguez-Alcayna M, Moya A, Baquero F, Latorre A. Complete genome sequence of *Acidaminococcus intestini* RYC-MR95, a Gram-negative bacterium from the phylum Firmicutes. J Bacteriol. 2011;193:7008–9.

37. Eribe ERK, Olsen I. *Leptotrichia* species in human infections II. J Oral Microbiol. 2017;9:1368848. doi: 10.1080/20002297.2017.1368848.

38. Sharma A. Virulence mechanisms of *Tannerella forsythia*. Periodontol 2000. 2010;54:106–16.

39. Hamlet SM, Taiyeb-Ali TB, Cullinan MP, Westerman B, Palmer JE, Seymour GJ. *Tannerella forsythensis* prtH genotype and association with periodontal status. J Periodontol. 2007;78:344–50.

40. Schisler TM, Bhavsar AK, Whitcomb BP, Freeman JH, Washington MA, Blythe JW, et al. Human papillomavirus genotypes in Pacific Islander cervical cancer patients. Gynecol Oncol Rep. 2018;24:83–6.

41. de Sanjose S, Quint WG, Alemany L, Geraets DT, Klaustermeier JE, Lloveras B, et al. Human papillomavirus genotype attribution in invasive cervical cancer: a retrospective cross-sectional worldwide study. Lancet Oncol. 2010;11:1048–56.

42. Cubie HA. Diseases associated with human papillomavirus infection. Virology. 2013;445:21–34.

43. Zhang Y, Xu X, Yu L, Shi X, Min M, Xiong L, et al. Vaginal microbiota changes caused by HPV infection in chinese women. Front Cell Infect Microbiol. 2022;12:814668. doi: 10.3389/fcimb.2022.814668.

44. Lin W, Zhang Q, Chen Y, Dong B, Xue H, Lei H, et al. Changes of the vaginal microbiota in HPV infection and cervical intraepithelial neoplasia: a cross-sectional analysis. Sci Rep. 2022;12:2812. doi: 10.1038/s41598-022-06731-5.

45. Chen C, Song X, Wei W, Zhong H, Dai J, Lan Z, et al. The microbiota continuum along the female reproductive tract and its relation to uterine-related diseases. Nat Commun. 2017;8:875. doi: 10.1038/s41467-017-00901-0.

46. Breshears LM, Edwards VL, Ravel J, Peterson ML. *Lactobacillus crispatus* inhibits growth of *Gardnerella vaginali*s and *Neisseria gonorrhoeae* on a porcine vaginal mucosa model. BMC Microbiol. 2015;15:276. doi: 10.1186/s12866-015-0608-0.

47. Fuochi V, Cardile V, Petronio Petronio G, Furneri PM. Biological properties and production of bacteriocins-like-inhibitory substances by *Lactobacillus* sp. strains from human vagina. J Appl Microbiol. 2019;126:1541–50.

48. Nunn KL, Wang Y-Y, Harit D, Humphrys MS, Ma B, Cone R, et al. Enhanced trapping of HIV-1 by human cervicovaginal mucus is associated with *Lactobacillus crispatus*-dominant microbiota. MBio. 2015;6:e01084–15.

49. Ceccarani C, Foschi C, Parolin C, D’Antuono A, Gaspari V, Consolandi C, et al. Diversity of vaginal microbiome and metabolome during genital infections. Sci Rep. 2019;9:14095. doi: 10.1038/s41598-019-50410-x.

50. Zhang Z, Li T, Zhang D, Zong X, Bai H, Bi H, et al. Distinction between vaginal and cervical microbiota in high-risk human papilloma virus-infected women in China. BMC Microbiol. 2021;21:90. doi: 10.1186/s12866-021-02152-y.

51. France MT, Mendes-Soares H, Forney LJ. Genomic comparisons of *Lactobacillus crispatus* and *Lactobacillus iners* reveal potential ecological drivers of community composition in the vagina. Appl Environ Microbiol. 2016;82:7063–73. doi: 10.1128/AEM.02385-16.

52. van Houdt R, Ma B, Bruisten SM, Speksnijder AGCL, Ravel J, de Vries HJC. *Lactobacillus iners*-dominated vaginal microbiota is associated with increased susceptibility to *Chlamydia trachomatis* infection in Dutch women: a case-control study. Sex Transm Infect. 2018;94:117–23.

53. Lennard K, Dabee S, Barnabas SL, Havyarimana E, Blakney A, Jaumdally SZ, et al. Microbial composition predicts genital tract inflammation and persistent bacterial vaginosis in South African adolescent females. Infect Immun. 2018;86. doi:10.1128/IAI.00410-17.

54. Molenaar MC, Singer M, Ouburg S. The two-sided role of the vaginal microbiome in *Chlamydia trachomatis* and *Mycoplasma genitalium* pathogenesis. J Reprod Immunol. 2018;130:11–7.

55. Cheong HC, Yap PSX, Chong CW, Cheok YY, Lee CYQ, Tan GMY, et al. Diversity of endocervical microbiota associated with genital *Chlamydia trachomatis* infection and infertility among women visiting obstetrics and gynecology clinics in Malaysia. PLoS One. 2019;14:e0224658. doi: 10.1371/journal.pone.0224658.

56. Filardo S, Di Pietro M, Tranquilli G, Sessa R. Biofilm in genital ecosystem: A potential risk factor for *Chlamydia trachomatis* infection. Can J Infect Dis Med Microbiol. 2019;2019:1672109.

57. Pekmezovic M, Mogavero S, Naglik JR, Hube B. Host-Pathogen interactions during female genital tract infections. Trends Microbiol. 2019;27:982–96.

58. BBMap. SourceForge. 2022. https://urldefense.com/v3/__https://sourceforge.net/projects/bbmap/__;!!LQC6Cpwp!oqX74jkqrDxLsVOjQjjwDwVAkoBqRkPDLTvAmDhQgFhfRZqnDnm-wrwbasbJWHztH0Ie_AOEBK8P_tCUTI9kDnd83np0BQ$. Accessed 2 Apr 2023.

59. Beghini F, McIver LJ, Blanco-Míguez A, Dubois L, Asnicar F, Maharjan S, et al. Integrating taxonomic, functional, and strain-level profiling of diverse microbial communities with bioBakery 3. Elife. 2021;10. doi:10.7554/eLife.65088.

60. CQLS Software Support. Metaphlan 3.0.14. 2022 https://software.cqls.oregonstate.edu/updates/metaphlan-3.0.14/. Accessed 2 Apr 2023.

61. Babiker A, Bradley HL, Stittleburg VD, Ingersoll JM, Key A, Kraft CS, et al. Metagenomic sequencing to detect respiratory viruses in persons under investigation for COVID-19. J Clin Microbiol. 2020;59. doi:10.1128/JCM.02142-20. Accessed 2 Apr 2023.

62. R: a language and environment for statistical computing. https://www.gbif.org/tool/81287/r-a-language-and-environment-for-statistical-computing. Accessed 2 Apr 2023.

63. Community Ecology Package [R package vegan version 2.6-4]. Comprehensive R Archive Network (CRAN); 2022. https://CRAN.R-project.org/package=vegan. Accessed 2 Apr 2023.

64. Adespatial: Multivariate Multiscale Spatial Analysis. Comprehensive R Archive Network (CRAN). https://cran.r-project.org/web/packages/adespatial/index.html. Accessed 2 Apr 2023.

65. Wickham H, Grolemund G. R for Data Science: Import, Tidy, Transform, Visualize, and Model Data. 2017. https://r4ds.had.co.nz. Accessed 2 Apr 2023.

66. PlotXTabs: Plot a Cross Tabulation of two variables using dplyr in CGPfunctions: Powell Miscellaneous Functions for Teaching and Learning Statistics. 2020. https://rdrr.io/cran/CGPfunctions/man/PlotXTabs.html. Accessed 2 Apr 2023.

67. R package FSA version 0.9.4. Comprehensive R Archive Network (CRAN); 2023. https://CRAN.R-project.org/package=FSA. Accessed 2 Apr 2023.

68. Kassambara A. ggpubr: “ggplot2” Based Publication Ready Plots. Github. https://github.com/kassambara/ggpubr. Accessed 2 Apr 2023.

69. Harrell FE Jr. Harrell Miscellaneous. R package Hmisc version 5.0-1. Comprehensive R Archive Network (CRAN); 2023. https://cran.r-project.org/web/packages/Hmisc/index.html. Accessed 2 Apr 2023.

70. Matrix v1.4 release. Matrix.org. 2023. https://matrix.org/blog/2022/09/29/matrix-v-1-4-release. Accessed 2 Apr 2023.

71. Hu Y-J, Lane A, Satten GA. A rarefaction-based extension of the LDM for testing presence-absence associations in the microbiome. Bioinformatics. 2021;37:1652–7.

72. McArdle BH, Anderson MJ. Fitting multivariate models to community data: A comment on distance-based reduncancy analysis. Ecology 2001;82:290–297.

