## Supplementary Figures for "The unique microbial diversity, community composition and networks of Pacific Islander endocervical and vaginal microbiomes in the presence or absence of *Chlamydia trachomatis* infection using metagenomic shotgun sequencing"

### The unique microbial diversity, community composition and networks of endocervical and vaginal microbiomes of Pacific Islanders based on comparative metagenomics

Sankhya Bommana[a], Yi-Juan Hu[b], Mike Kama[c], Ruohong Wang[a], Reshma Kodimerla, Kenan Jijakli[d], Timothy D. Read[d*], Deborah Dean[a,e,f,g,h*]

[a]Department of Pediatrics, University of California San Francisco, Oakland, CA, USA
[b]Department of Biostatistics and Bioinformatics, Emory University, Atlanta, GA, USA
[c]Ministry of Health and Medical Services, Suva, Fiji
[d]Department of Medicine, Emory University School of Medicine, Atlanta, Georgia, USA
[e]Department of Medicine, University of California San Francisco, San Francisco, CA, USA
[f]Department of Bioengineering, Joint Graduate Program, University of California San Francisco and University of California Berkeley, San Francisco, CA, USA
[g]Bixby Center for Global Reproductive Health, University of California San Francisco, San Francisco, CA, USA
[h]Benioff Center for Microbiome Medicine, University of California San Francisco, San Francisco, CA, USA

**Supplementary Table 7**. Accession numbers for all metagenomic samples submitted to NCB.

Supplementary Figure 1

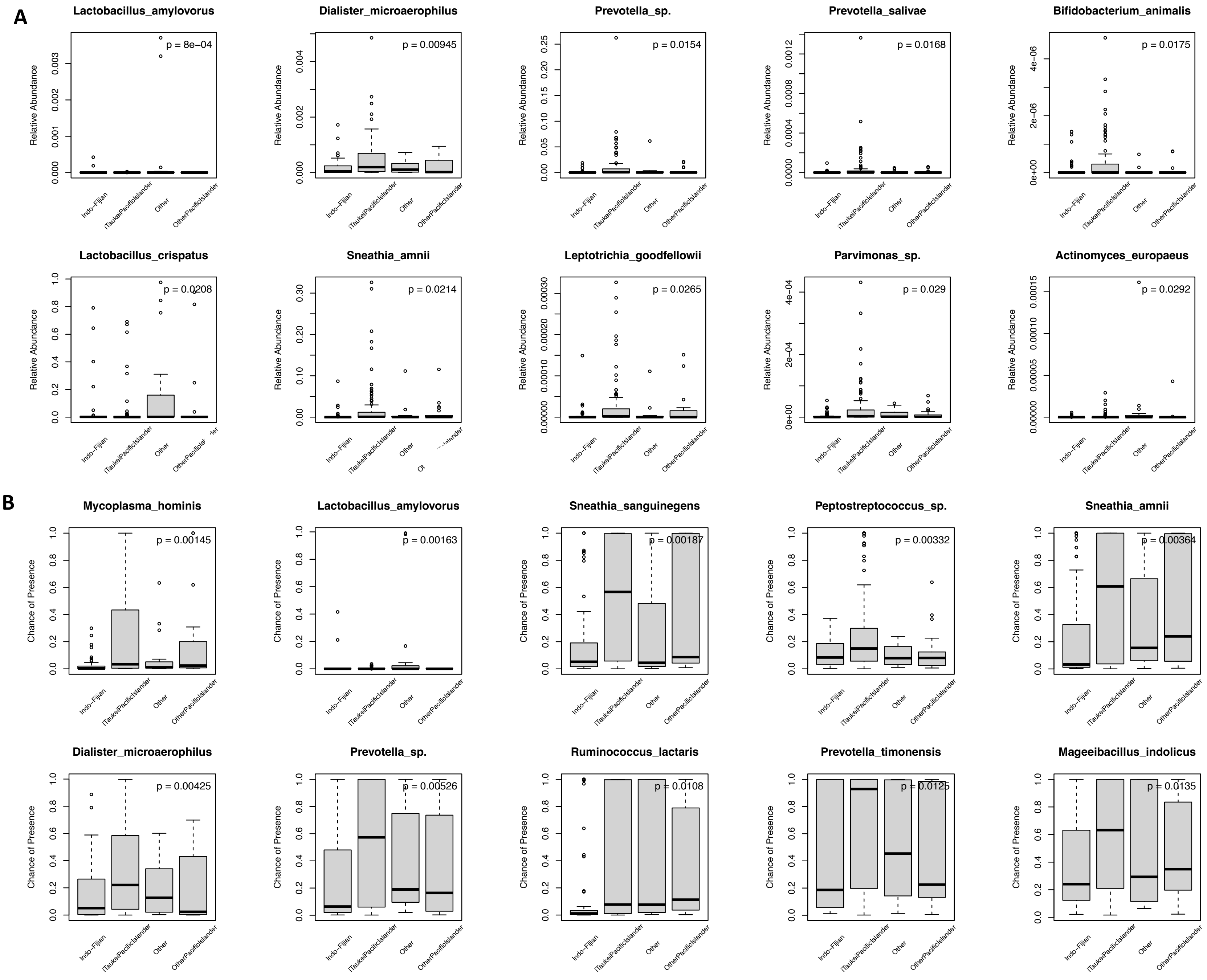

Supplementary Figure 2

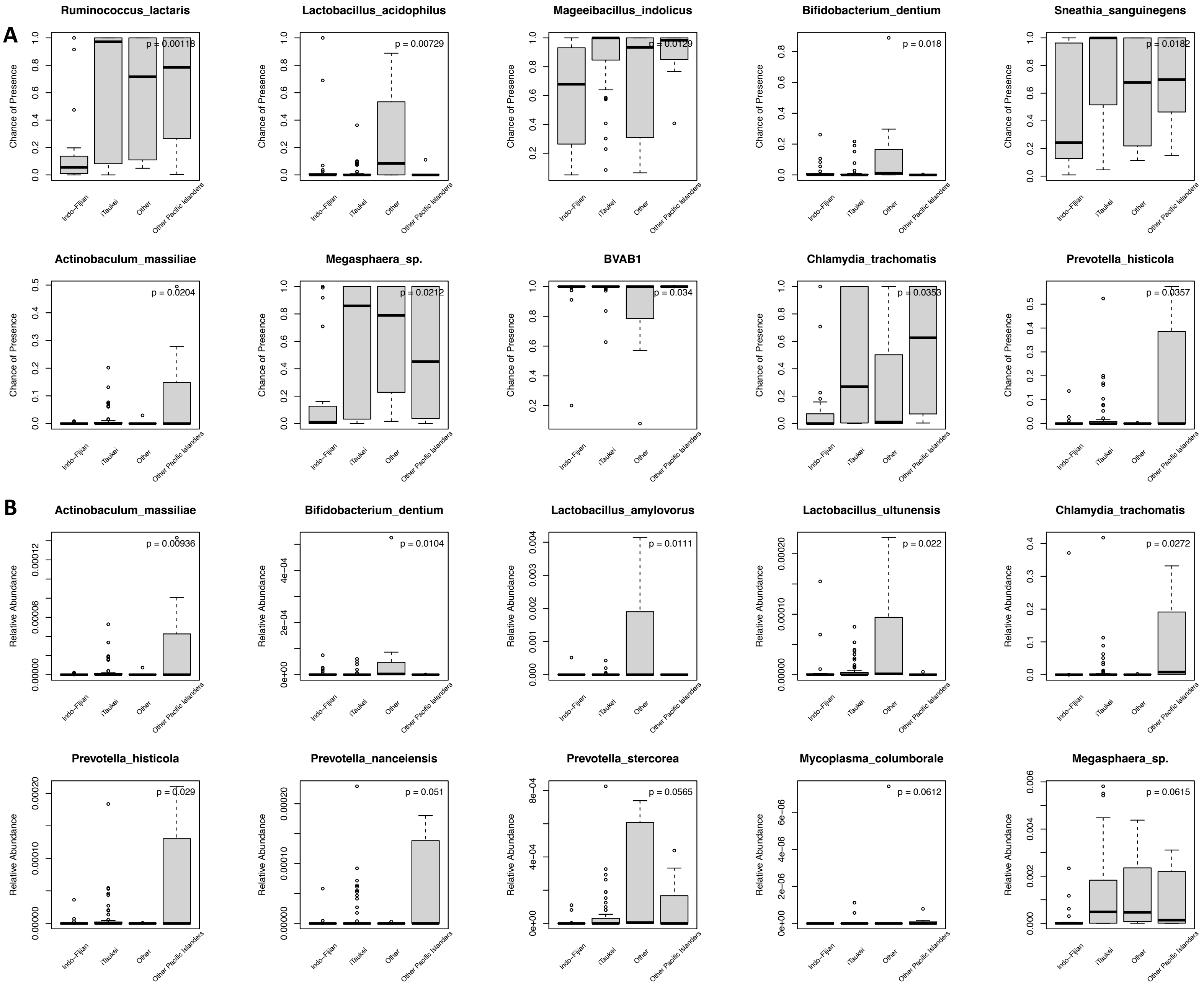

### Supplementary Figure 3A

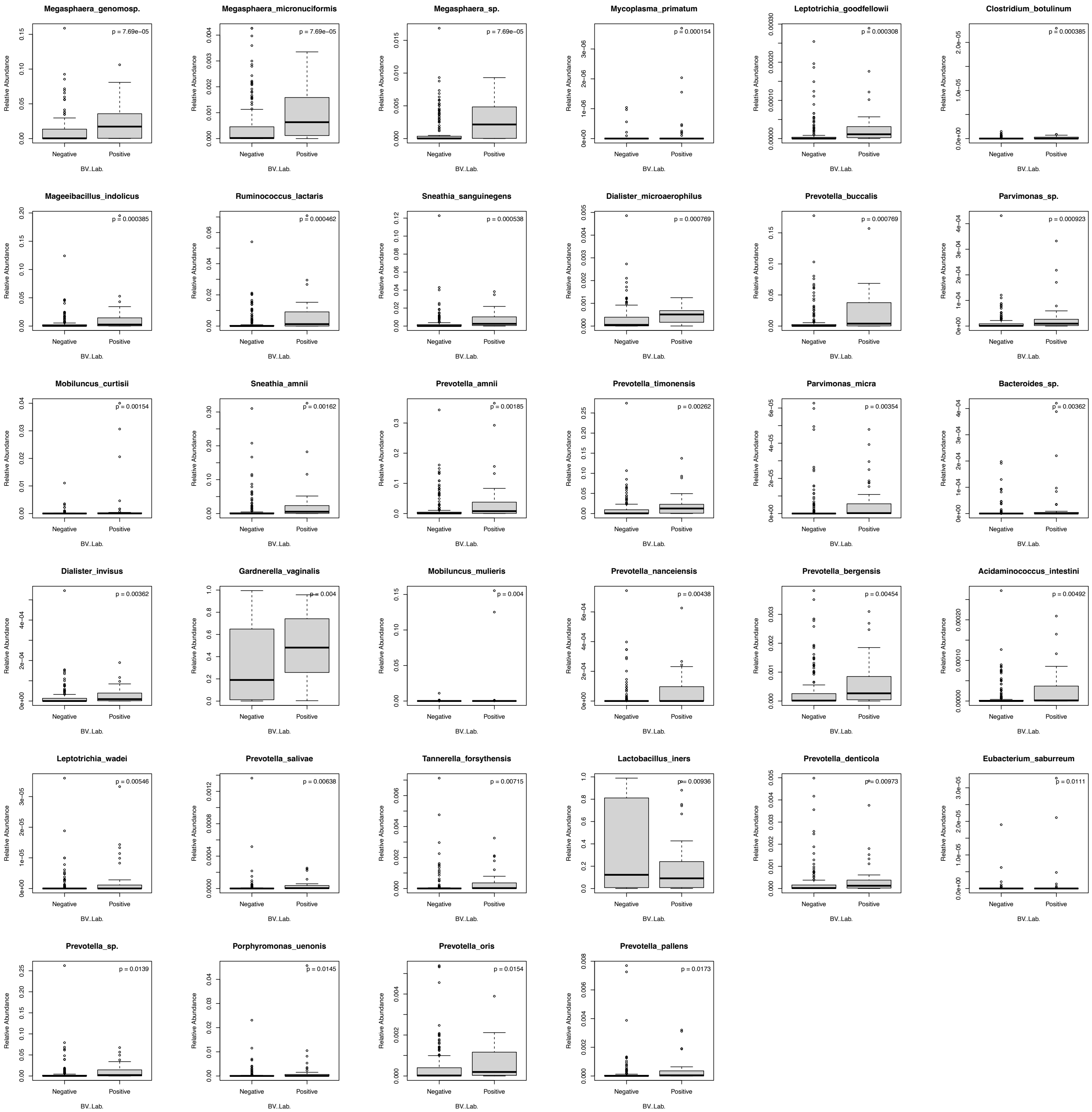

### Supplementary Figure 3B

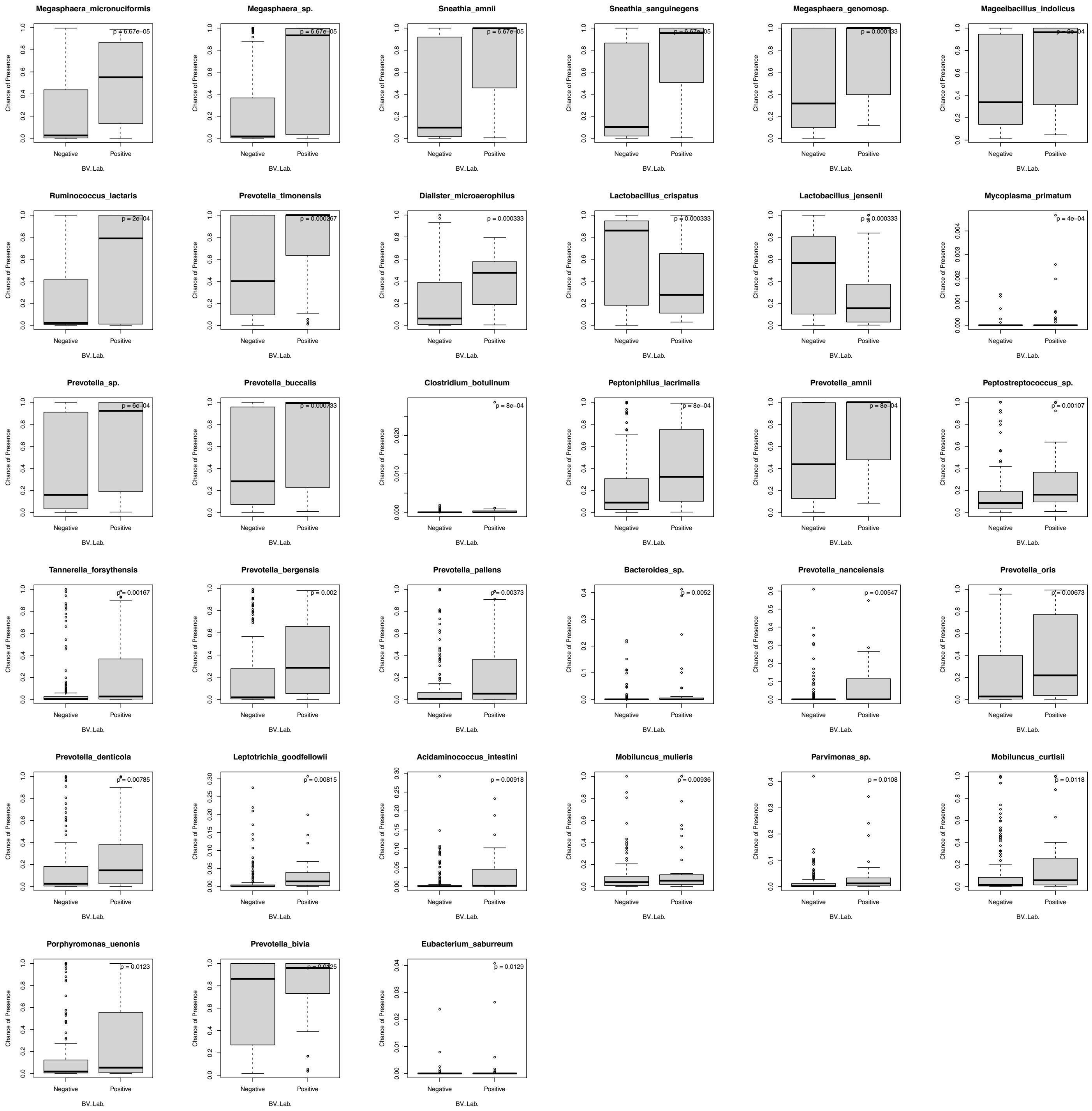

Supplementary Figure 4

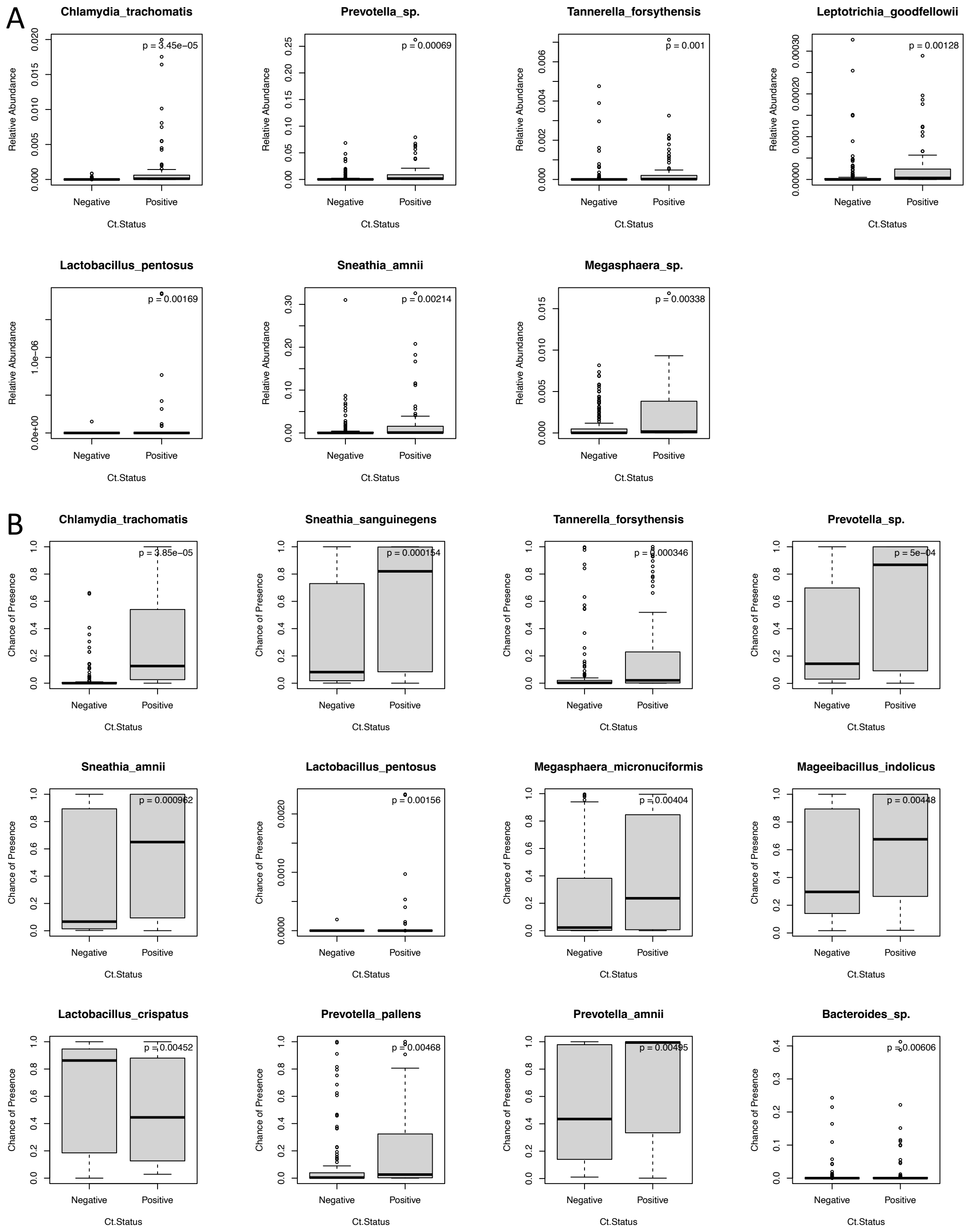

### Supplementary Figure 5

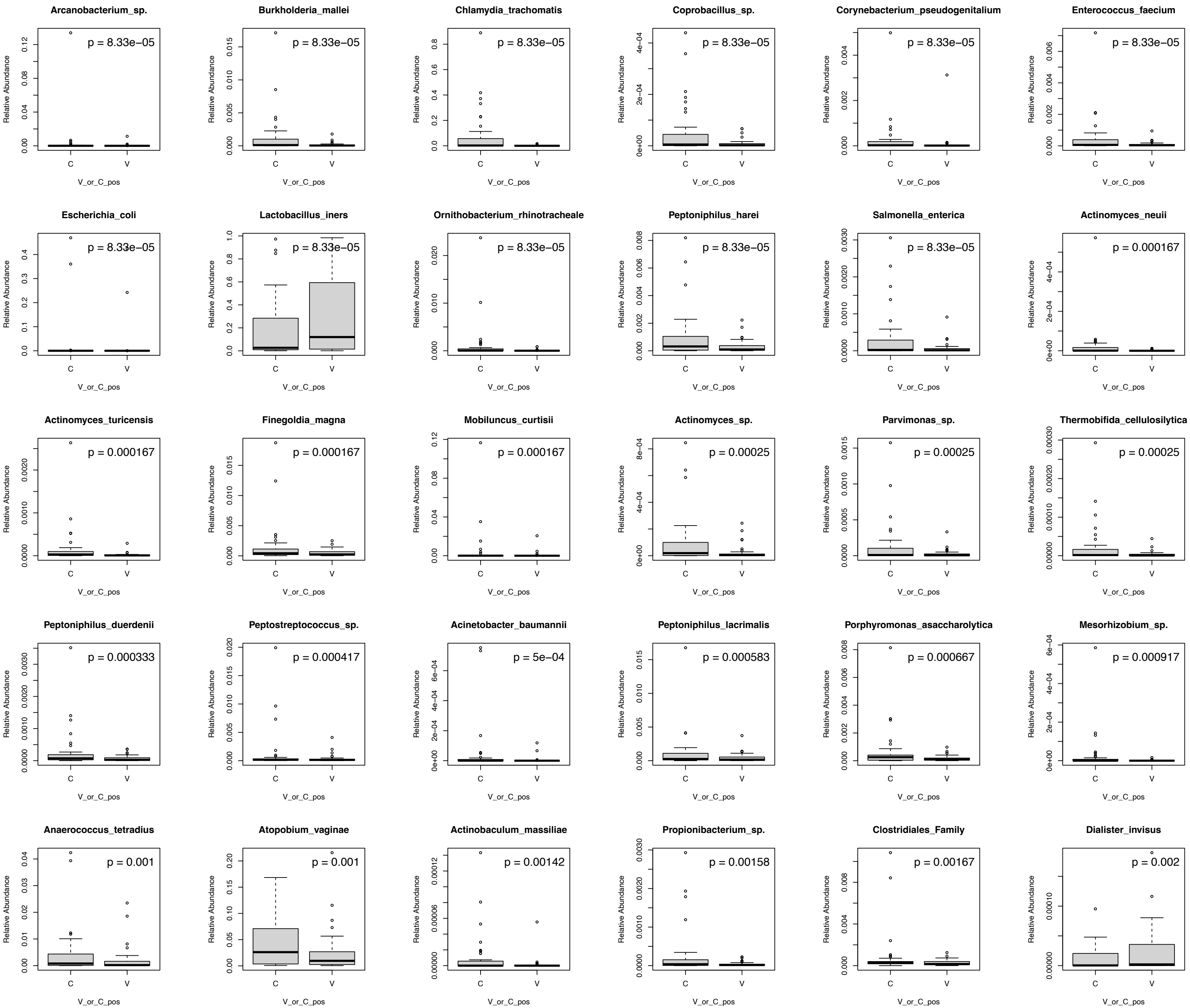

Supplementary Figure 5 (cont.)

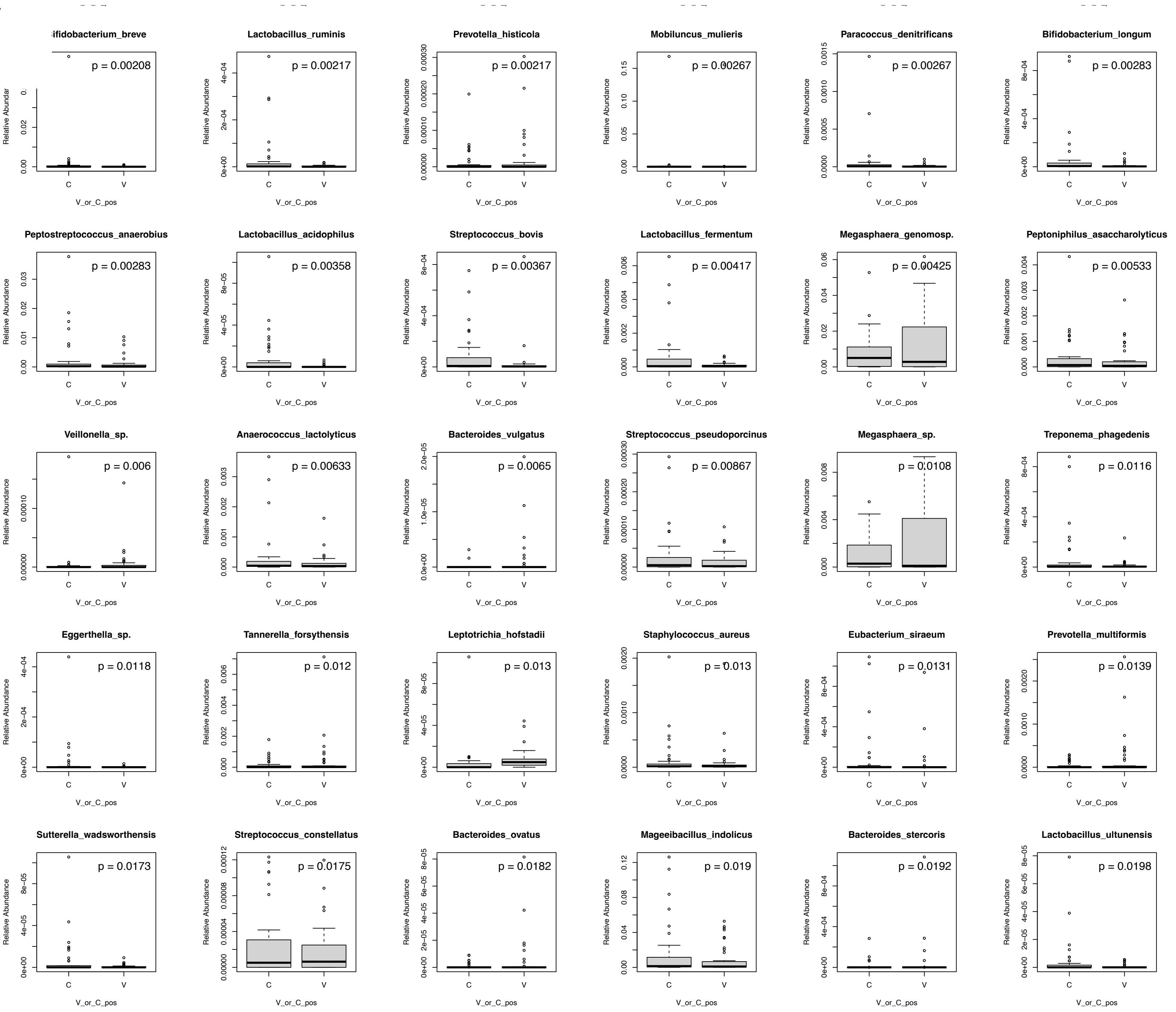

Supplementary Figure 5 (cont.)

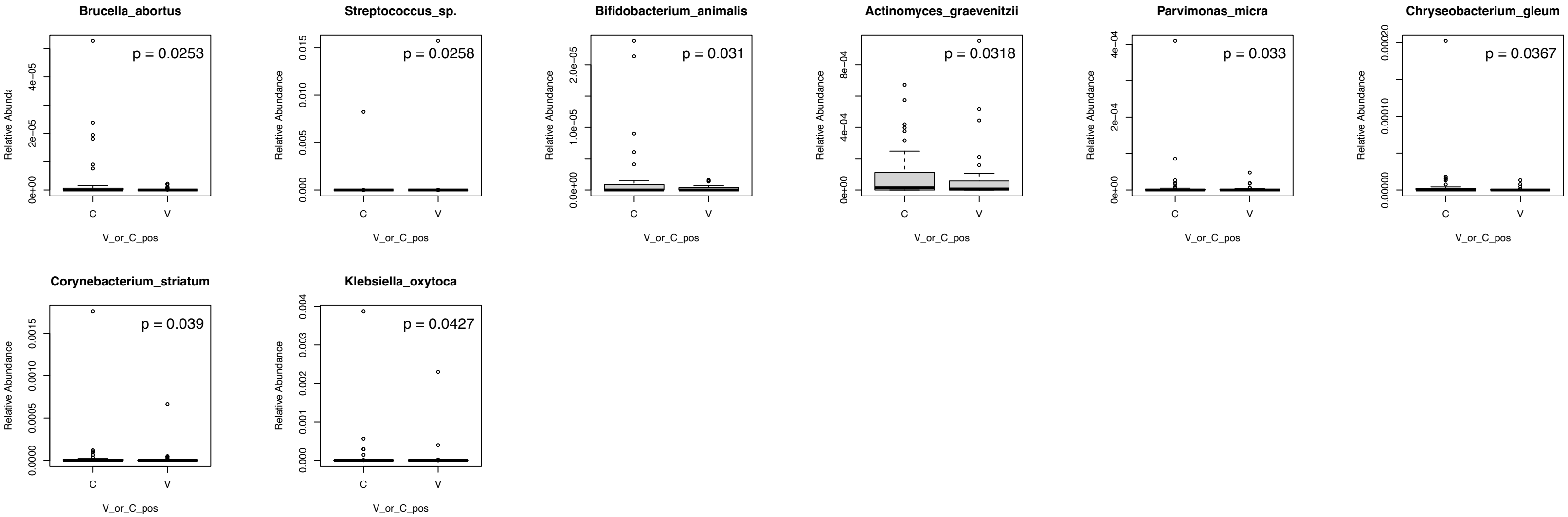

Supplementary Figure 6

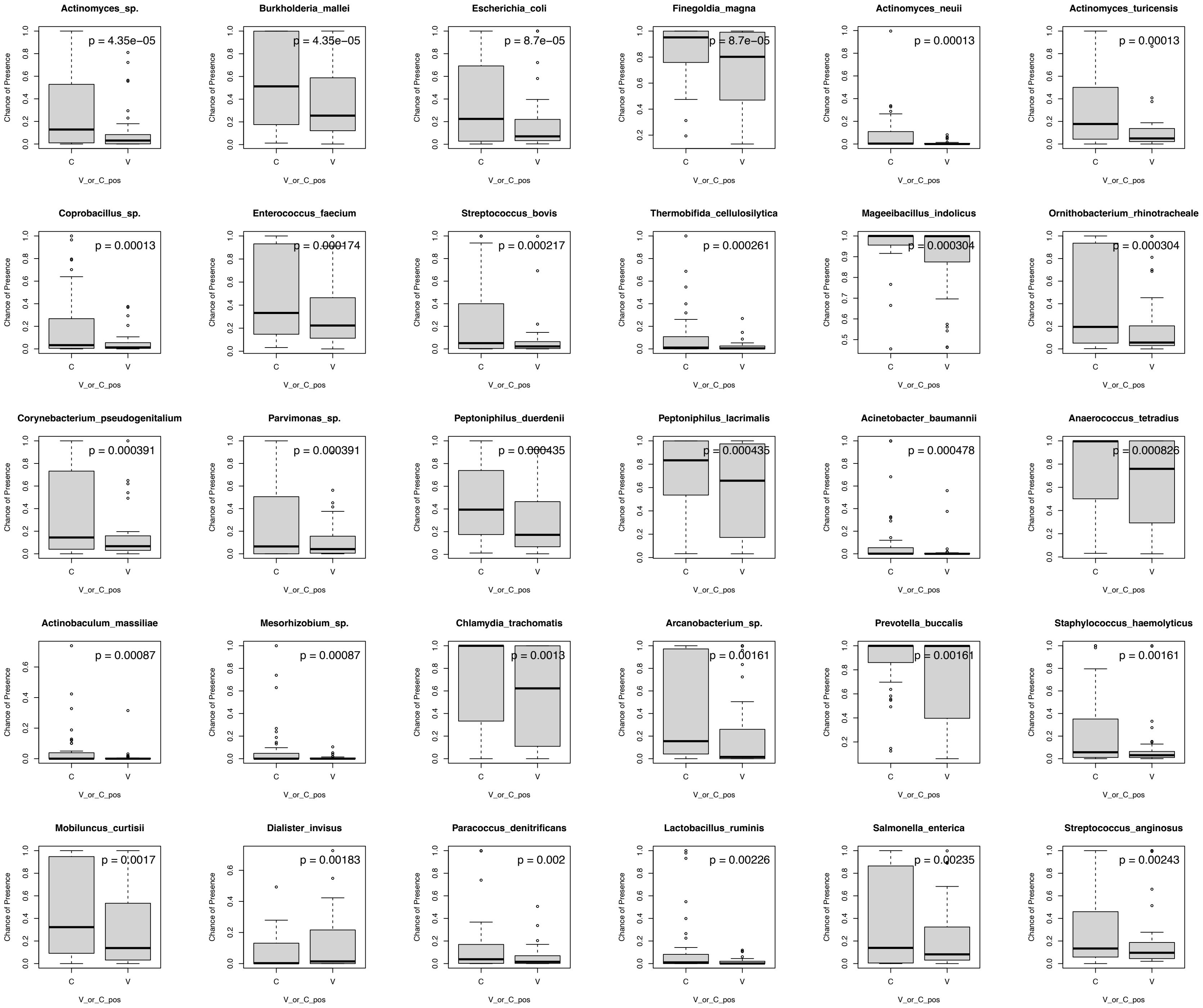

Supplementary Figure 6 (cont.)

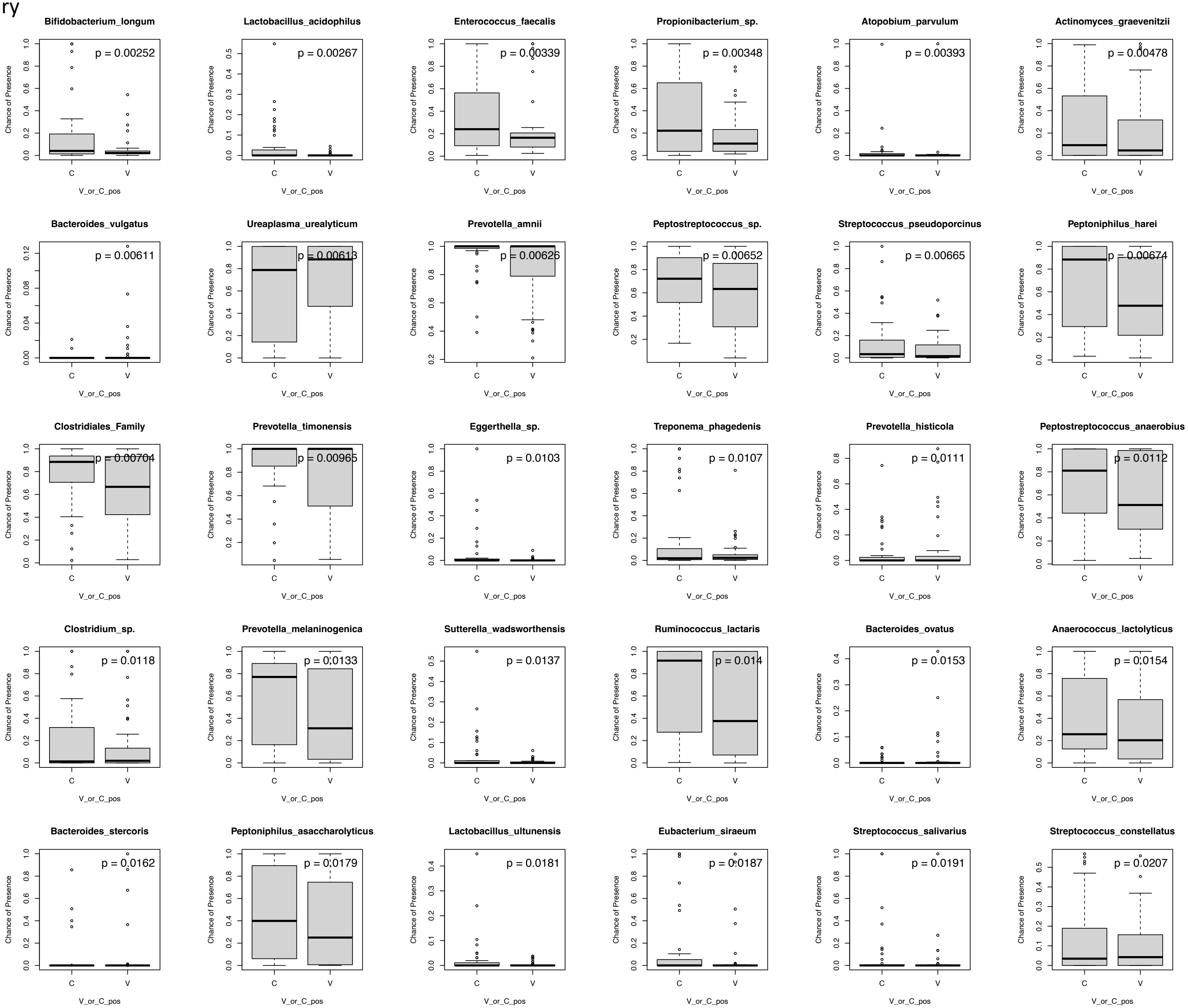

Supplementary Figure 6 (cont.)

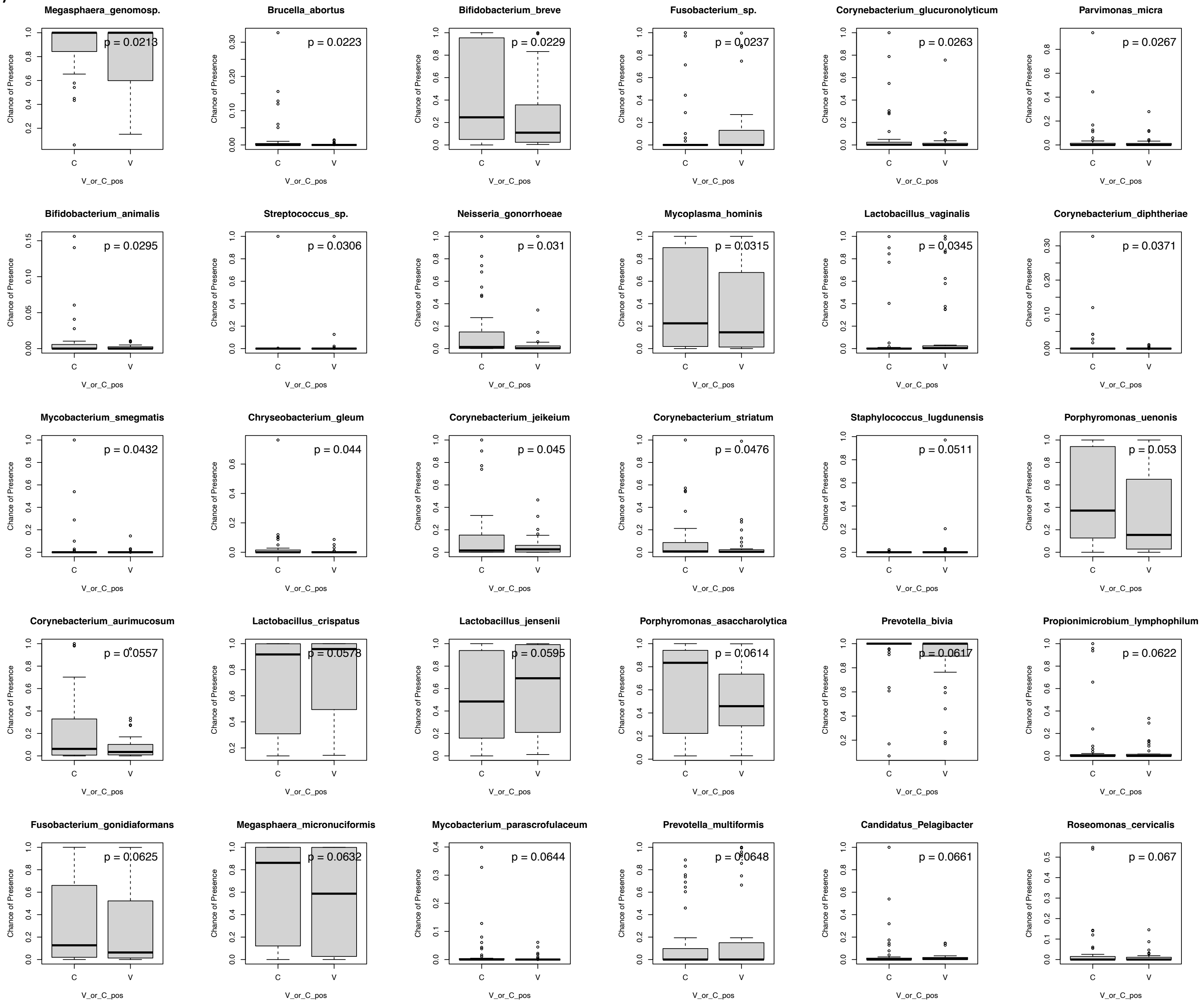

### Supplementary Figure 7

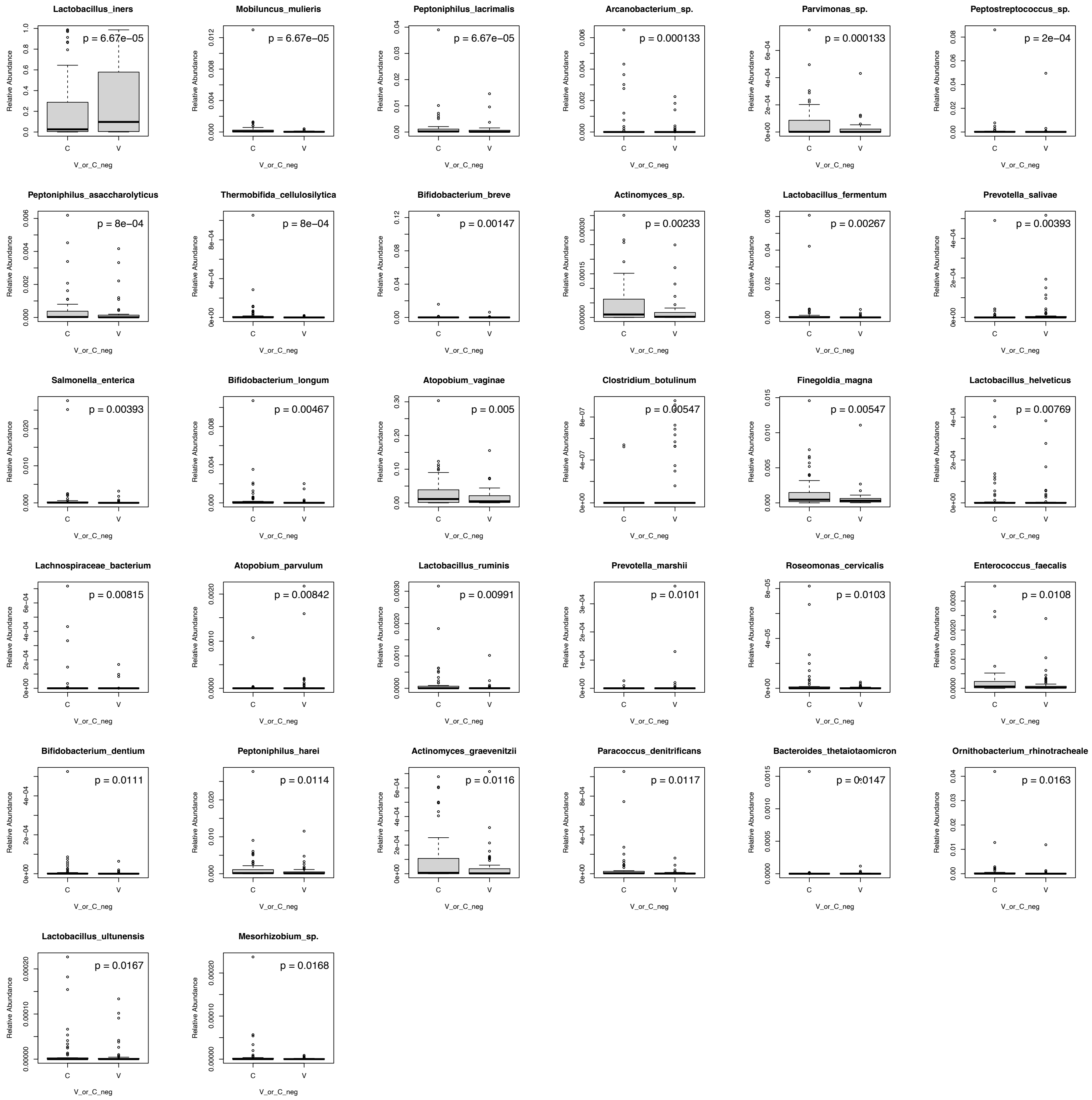

Supplementary Figure 8

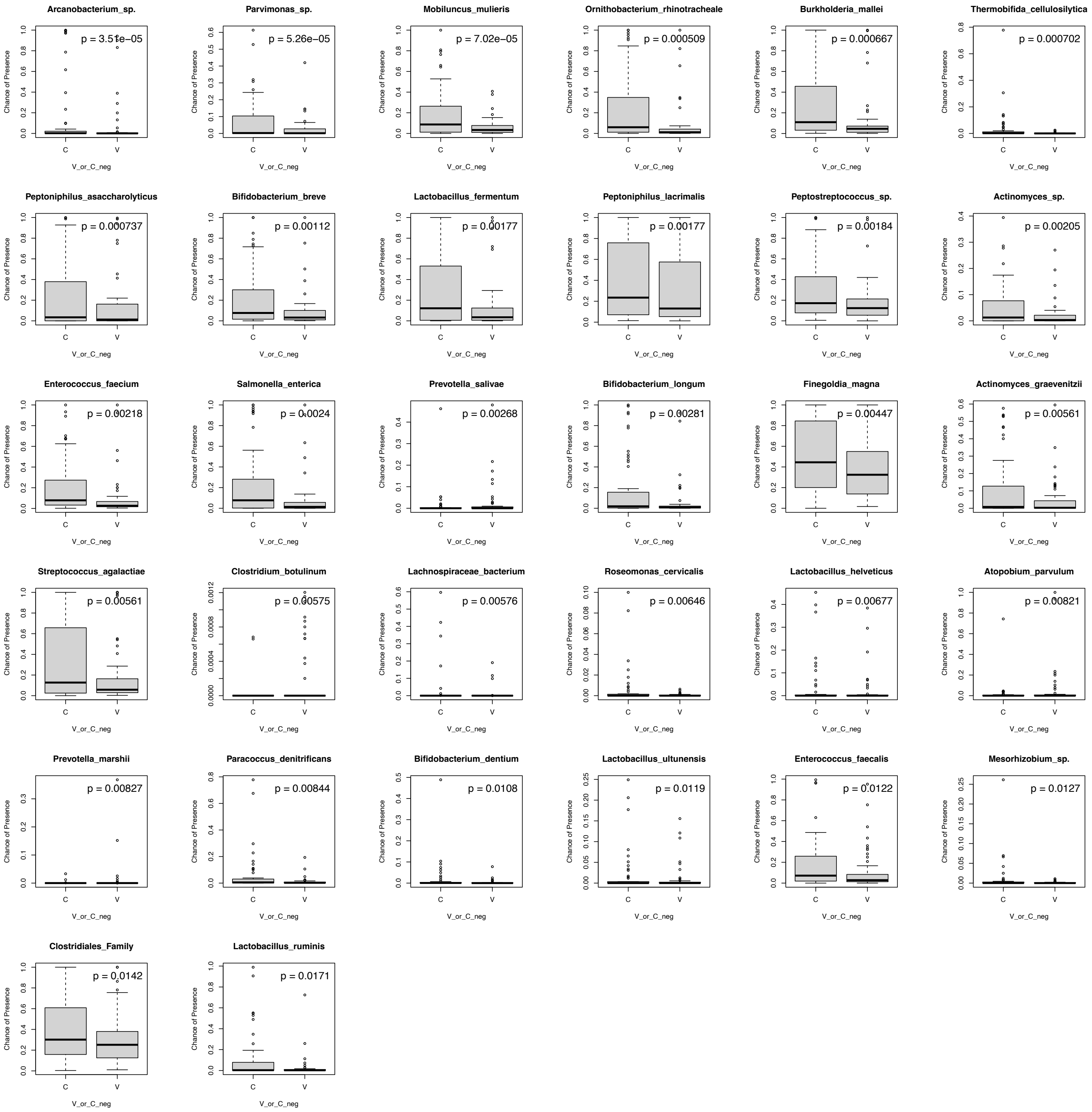

### Supplementary Figure 9

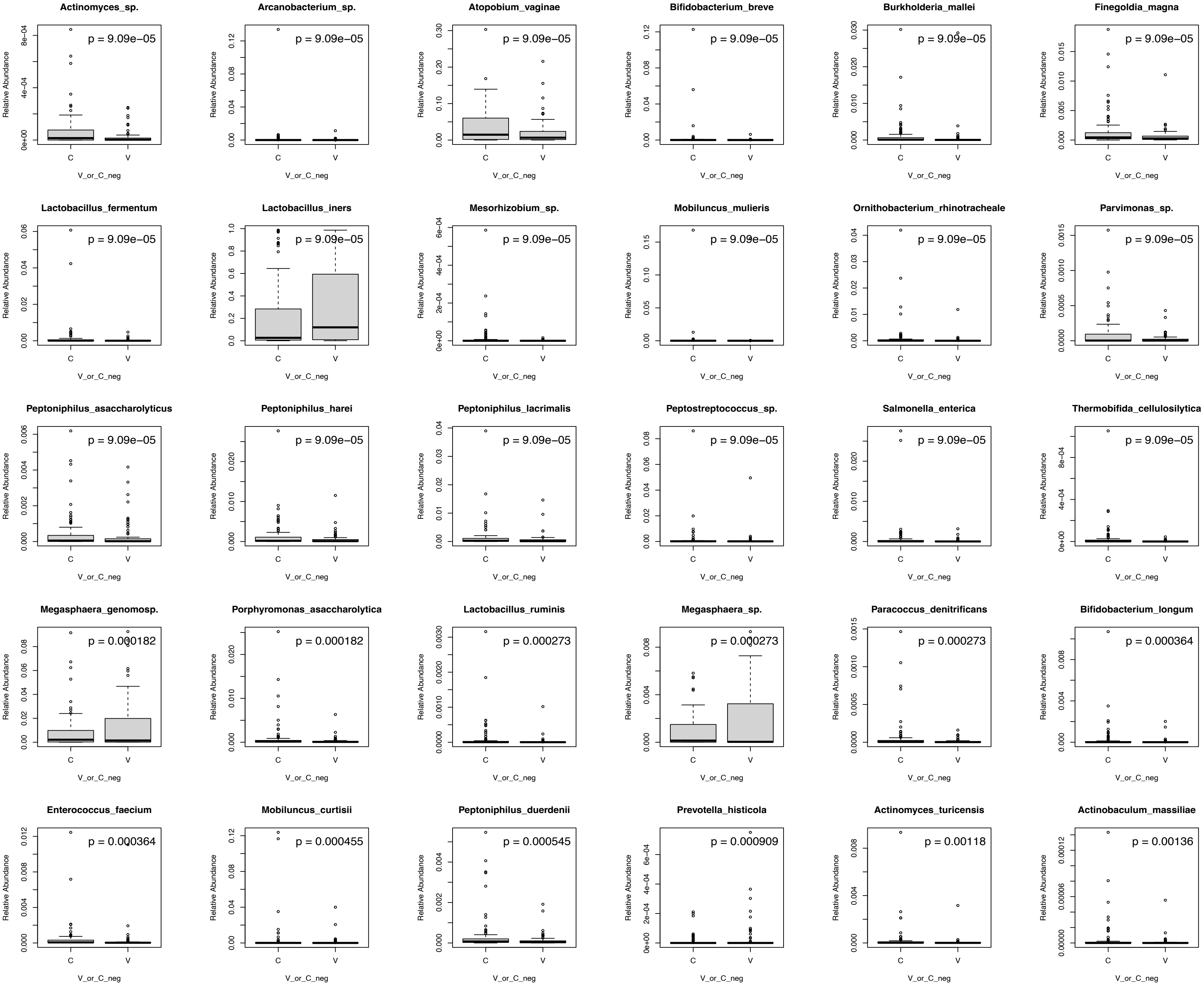

Supplementary Figure 9 (cont.)

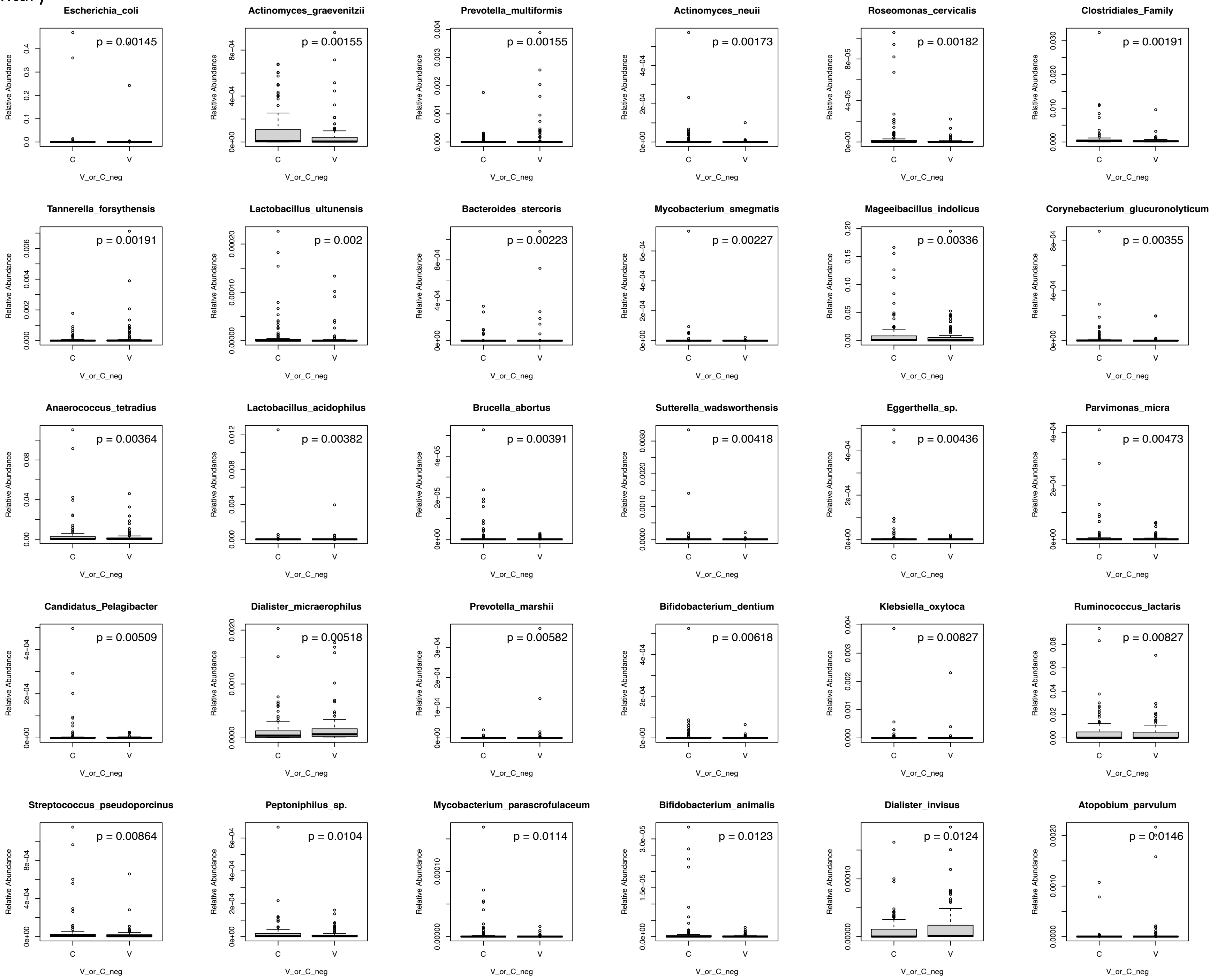

Supplementary Figure 9 (cont.)

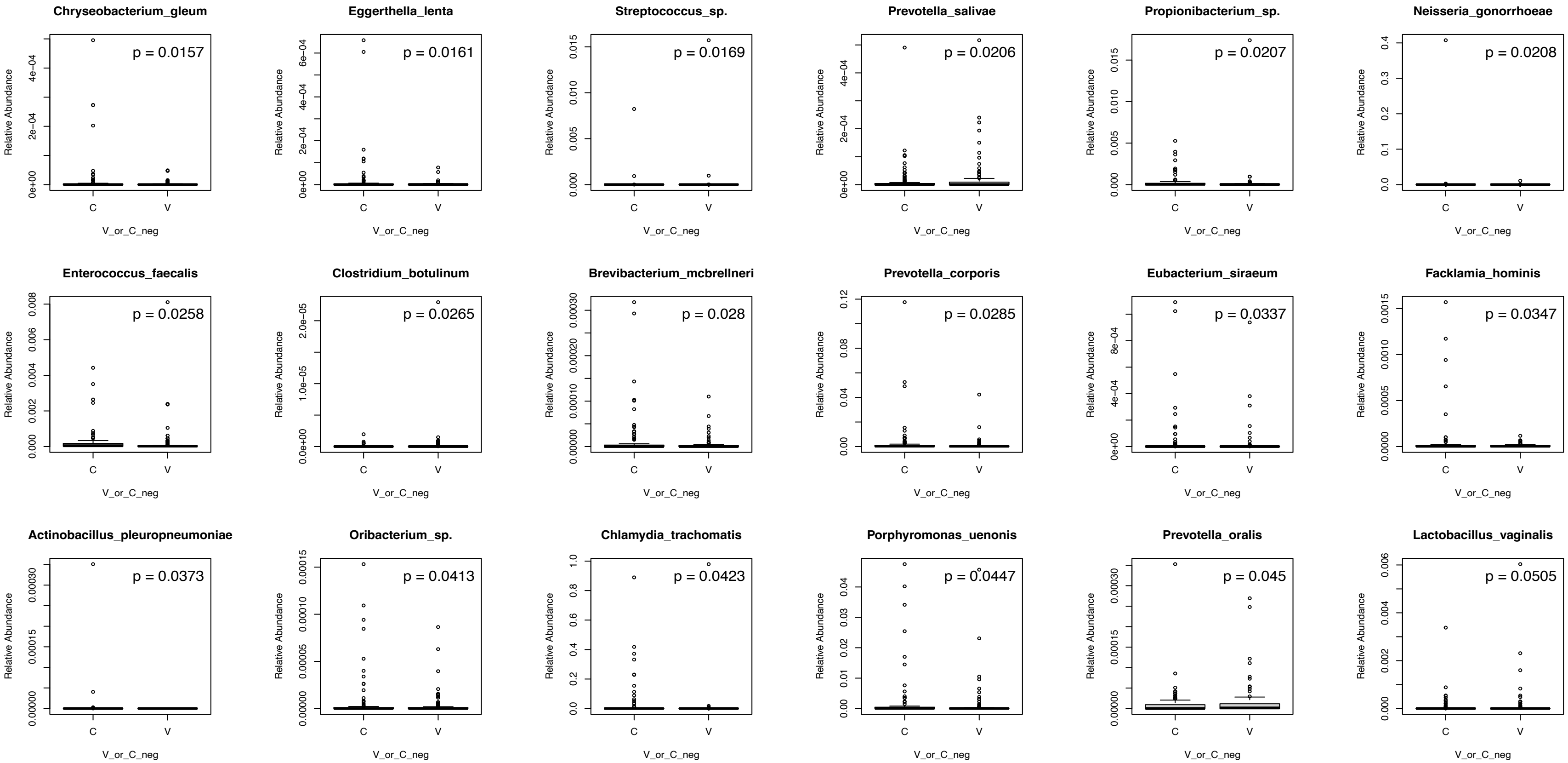

Supplementary Figure 10

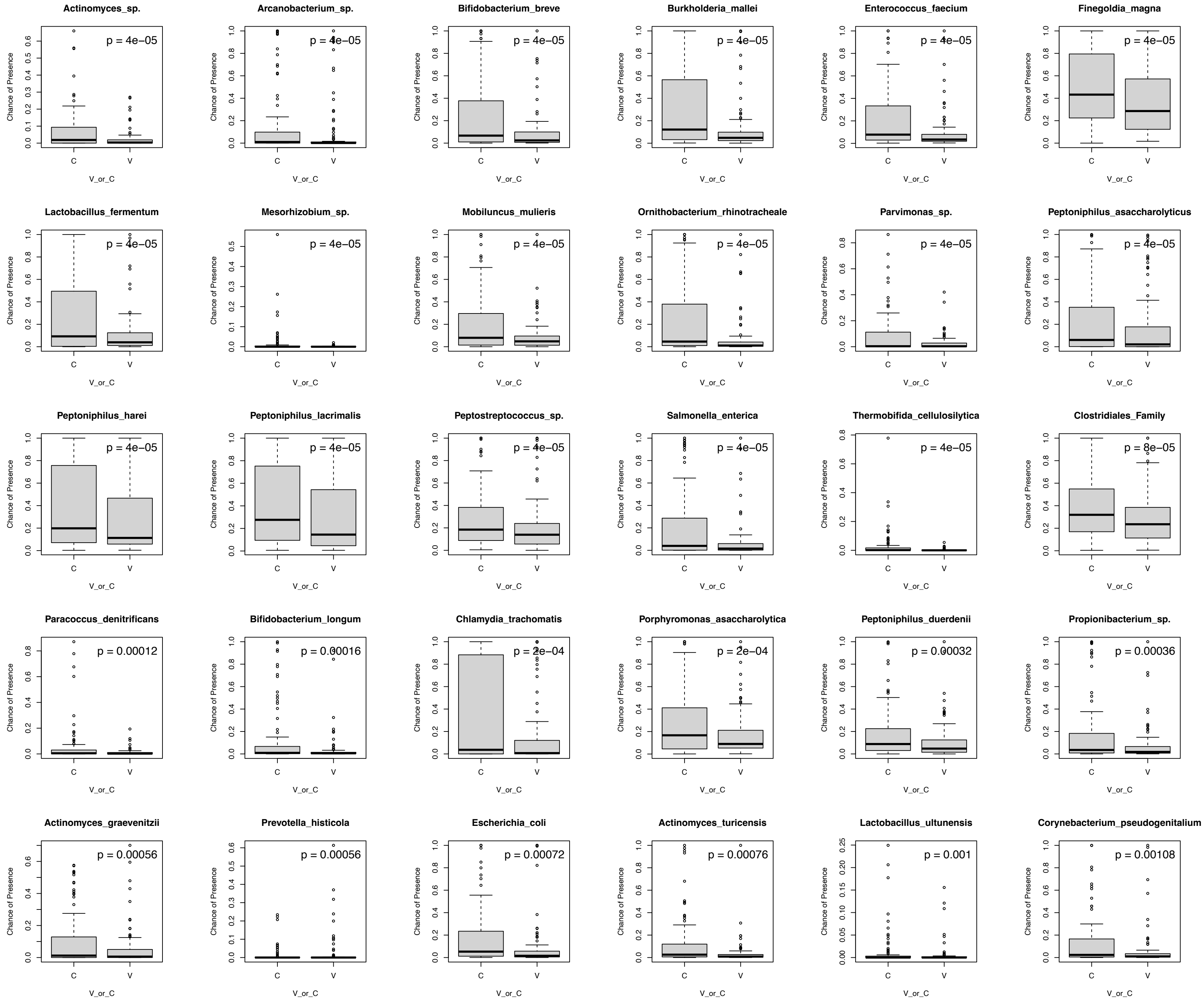

Supplementary Figure 10 (cont.)

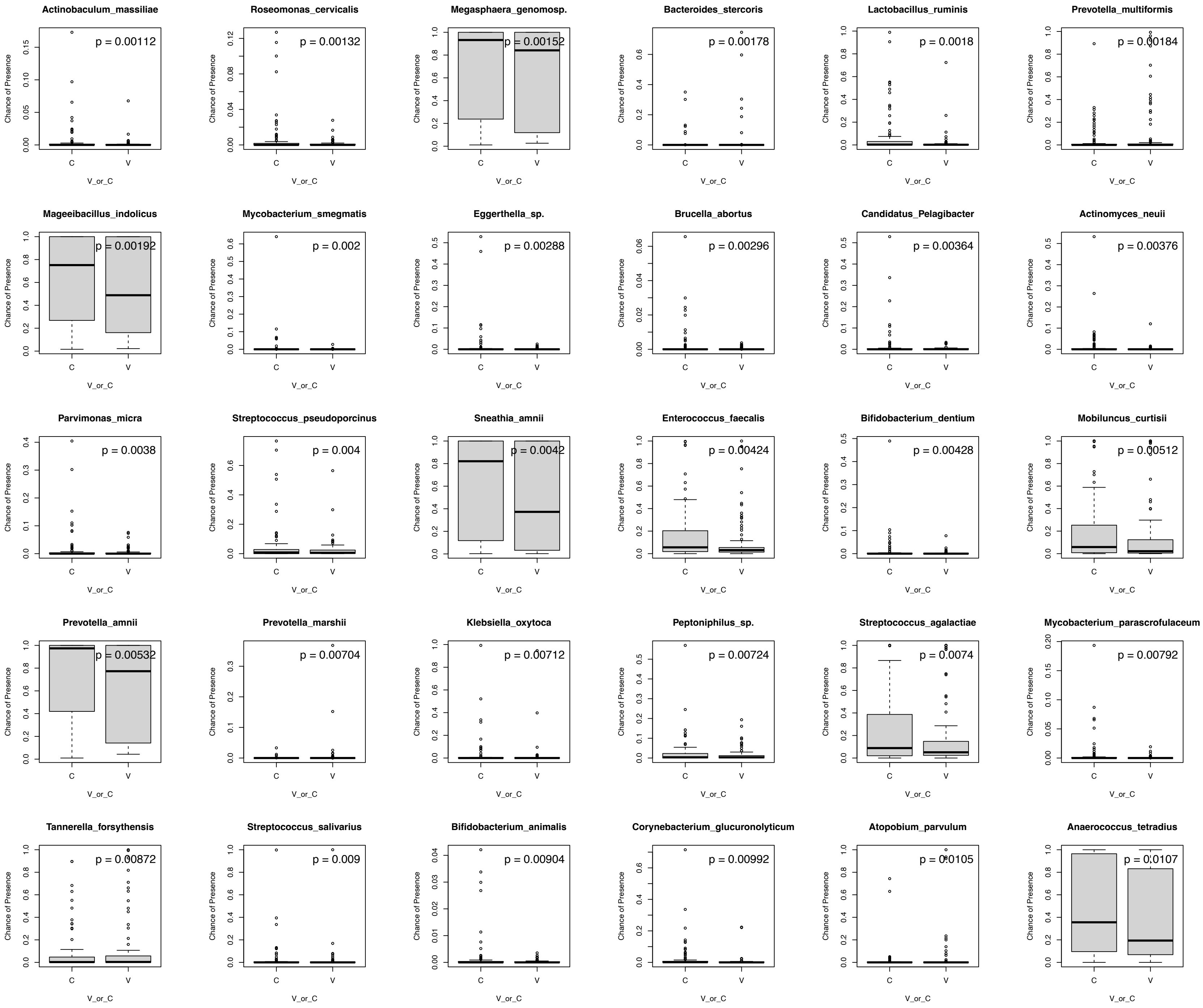

Supplementary Figure 10 (cont.)

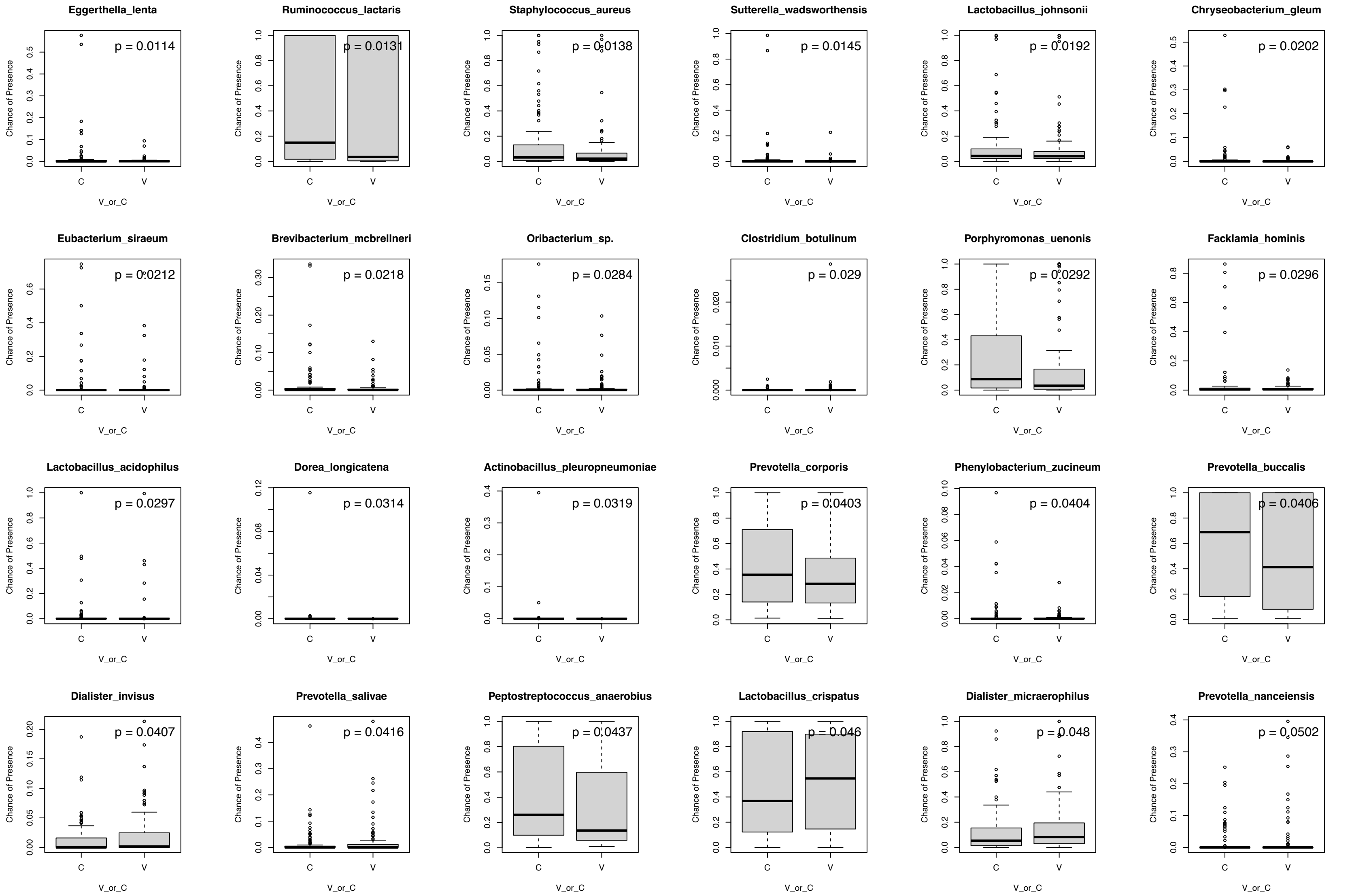
